# Spatially confined niches support hypoxia-associated transcriptional plasticity contributing to malignant progression in IDH-mutant gliomas

**DOI:** 10.64898/2026.05.31.729105

**Authors:** Depro Das, Alina Pandele, Vatsal D. Jariwala, Nishita S. Ghariwala, Aso O. Mohammed, Tonmoy Das, Christopher Krolla, Saif Mukramoon Arosh, Naiem Hossain, Mohammad Al Shhab, Ismail Hosen, Stuart J. Smith, Jürgen Grauvogel, Ioannis Vasilikos, Roland Roelz, Julia Nakgawa, Rouven Hoefllin, Itay Tirosh, Tareq A. Juratli, Matthias Schneider, Marco Prinz, Shawn Hervey Jumper, Ruman Rahman, Melanie Boerries, Jürgen Beck, Kevin Joseph, Geoffroy Andrieux, Vidhya M. Ravi, Sajib Chakraborty

## Abstract

**Background:** While hypoxia is a well-established driver of glioblastoma progression, its role in IDH-mutant gliomas, characterized by localized hypoxic microenvironments rather than overt necrosis, remains poorly understood. Here, we investigate how hypoxia and microenvironmental-adaptations shape cellular heterogeneity and transcriptional plasticity in these tumors.

**Methods:** We integrated bulk, single-cell, and spatial-transcriptomics datasets from IDH-mutant glioma patients (Astrocytomas and Oligodendrogliomas) to characterize cellular-states and map the localization of hypoxic niches. To uncover tumor microenvironment effects, we established co-culture models using primary IDH-mutant glioma cells with human microglia and astrocytes, maintained under hypoxic and normoxic conditions, followed by bulk RNA-sequencing.

**Results:** We identified a hypoxia-associated astrocyte-like (AC-like) program that defines a quiescent, non-cycling population with a distinct transcription factor profile indicative of functional plasticity in IDH-mutant gliomas. These cells harbor glioma stem cell (GSC)-like features and are poised for a quiescent-to-activated (Q-to-A) transition that drives tumor progression. Mechanistically, co-culture models reveal that microglia promote this Q-to-A transition by enhancing HBEGF/EGFR paracrine signaling. Spatial transcriptomics uncovers the co-localization of hypoxic niches within quiescent AC-like cells, whereby the activated subpopulation forms discrete niches defined by localized HBEGF/EGFR communication gradients. Notably, tumors exhibiting elevated EGFR-driven activation signatures correlate with higher histological grade and poorer patient survival, implicating the Q-to-A transition as a critical driver of malignant progression.

**Conclusion:** Q-to-A transition within the hypoxic niche represents a critical driver of malignant progression in IDH-mutant gliomas, providing a microenvironment-driven mechanism for the transition to higher-grade disease and identifying targetable-vulnerabilities for therapeutic intervention.

**Key points:**

- Hypoxic niches spatially confine quiescent, astrocyte-like cellular state characterized by glioma stem cell features in IDH-mutant gliomas.
- Microglia triggers the quiescent-to-activated (Q-to-A) transition via paracrine EGFR signaling crosstalk.
- EGFR-driven Q-to-A plasticity serves as a microenvironmentally-regulated driver of malignant progression and adverse patient survival.

**Importance of the study:** While high-grade glioblastomas feature well-defined hypoxic and necrotic regions that drive tumor progression and therapy resistance, IDH-mutant gliomas lack these distinct hallmarks, instead exhibiting disorganized hypoxic niches. Consequently, the functional contribution of these niches to tumor progression has remained poorly understood. Although genetic and epigenetic alterations are established drivers of progression in IDH-mutant gliomas, we provide an additional, essential layer of complexity: microenvironment-driven transcriptional plasticity. By uncovering the relationship between tumor hypoxia and heterogeneous glioma cell states, we uncover that these niches are essential for such plasticity. We also show that the tumor microenvironment provides critical molecular cues necessary to initiate a transcriptional shift enabling glioma cells to transition from a quiescent reservoir into an activated state primed for rapid proliferation. By identifying this paracrine mechanism, our work uncovers a targetable vulnerability that dictates how dormant cell reservoirs are mobilized to fuel malignant transformation.

## Introduction

Hypoxia, a hallmark of the glioma tumor microenvironment (TME), results from the imbalance between oxygen supply and demand. While physiological oxygen tension in adult human tissues ranges from 2-9% O_2_, oxygen levels within glioma tumor cores and peritumoral regions drop to ∼1.25% and ∼2.5%, respectively^1,2^. This chronic hypoxia initiates adaptive responses that foster aggressive tumor traits, including genomic instability, impaired DNA mismatch repair, and elevated mutational burden, ultimately driving tumor evolution and resistance to therapy^1,2^. Additionally, hypoxic signaling is intricately linked to tumorigenesis and it influences the outcome of cancer treatments by interacting with different cellular pathways. Central to this process are hypoxia-inducible factors (HIFs), transcriptional regulators stabilized under low oxygen conditions. HIFs promote tumor progression through angiogenesis, epithelial-mesenchymal transition (EMT), altered proliferation, and metabolic reprogramming^1,3^. In immune and stromal cells, HIF signaling regulates chemotaxis, phagocytosis, oxidative burst, and apoptosis, enabling adaptation to hypoxic stress. Specifically, HIF-1α has been linked to glioma invasiveness and pro-angiogenic factor expression^1^, whereas HIF-2α impairs inflammatory resolution and reduces treatment efficacy by inducing a dormant state in tumor-associated myeloid cells^2^. Thus, hypoxia not only affects tumor cells but also reshapes stromal/immune cell phenotypes within the glioma TME.

Intratumoral heterogeneity further amplifies this complexity. Glioma cells within the TME showcase diverse dynamic transcriptional cellular states and expression programs reflecting a putative cellular hierarchy. This cellular hierarchy reflects neural cell types which mimic the early development of a human brain. Such transcriptional states for IDH-mutant gliomas include a stem-like program that resembles neural progenitor cells (NPC-like), while astrocyte-like (AC-like) and oligodendrocyte-like (OC-like) programs resemble differentiated glial lineages^4^. Dynamic plasticity across these cellular states is a defining hallmark of malignant brain tumors, strongly influenced by hypoxia, and which substantially promotes transcriptional reprogramming. Recent advances now extend these glioma metaprograms into a unifying framework of activation state architecture (ASA), which resolves tumor cells along a continuum of quiescence (Q), activation (A), and differentiation (D)^5^. Notably, hypoxia plays a central role in modulating tumor hierarchies, with malignant programs partitioning into quiescent and activated stages. Such a framework emphasizes that cellular plasticity is governed by dynamic state transitions, with the quiescence-to-activation (Q-to-A) transition emerging as a critical determinant of tumor growth and therapy resistance^5^. Despite these advances, the interplay between transcriptional programs, cellular plasticity, and hypoxia in IDH-mutant gliomas remains largely unexplored.

In this study, we combined bulk, single-cell, and spatial transcriptomics with co-culture models to investigate the hypoxic microenvironment of IDH-mutant gliomas. We examined the association between hypoxia and glioma-specific transcriptional programs, with a particular focus on microenvironmental cues that influence cellular state plasticity. Our framework elucidates the mechanisms coupling hypoxia to adaptive glioma cell states and tumor heterogeneity.

## Methods

### Data acquisition and sample attributes

Single-cell RNA sequencing (scRNA-seq) datasets from two separate human glioma studies (Wang et al.,^6^ and Abdelfattah et al.,^7^) were obtained from NCBI GEO (GEO accessions: GSE138794 and GSE182109). We also retrieved eight IDH-mutant glioma samples profiled by 10x Visium spatial transcriptomics (stRNA-seq) from Greenwald et al.,^8^ (GSE237183) and He et al.,^9^ (GSE270355). For large-scale validation, the ‘Brain Lower Grade Glioma (TCGA, PanCancer Atlas)’ cohort, consisting of bulk RNA-seq, whole exome sequencing (WES), CNA, and DNA methylation data with associated clinical information, was downloaded from cBioPortal^10,11^. Patient-associated clinical information regarding age, gender, sample type, glioma grades, histologic classification, and overall survival was primarily focused upon in our study.

### scRNA-seq data preprocessing, integration, and annotation

Single-cell data specific for IDH-mutant gliomas from Wang et al.,^6^ and Abdelfattah et al.,^7^ were processed using ‘Seurat’ (v5.2.0). See details in Supplementary Methods.

### Non-negative matrix factorization (NMF) based metaprogram identification

To characterize cellular heterogeneity within the malignant compartment and identify robust transcriptional states, Non-negative Matrix Factorization (NMF) was applied. See details in Supplementary Methods.

### Gene signature scoring

Module scores for IDH-mutant glioma MPs and QAD states^5^ were computed using Seurat’s ‘AddModuleScore’ function, extended with curated gene sets from several landmark studies^12–17^ to profile cellular states comprehensively. See details in Supplementary Methods. See details in Supplementary Methods.

### Pathway enrichment and transcription factor activity inference

PROGENy model was applied to infer pathway activities and transcription factor (TF) activities were inferred using the CollecTRI regulatory network^18^, implemented through the ‘decoupleR’^19^. See details in Supplementary Methods.

### Pseudotime trajectory analysis

See details in Supplementary Methods.

### Cell-cell interaction analysis in scRNA-seq data

We utilized ‘CellChat’^20^ and ‘LIANA’^21^ to infer and analyze cell-cell communication (CCC) networks between glioma and immune cell populations. See details in Supplementary Methods.

### Cell co-culture and hypoxia treatment

A glioma-glial co-culture system was established using patient-derived IDH-mutant glioma cells (UKFR #971), human microglia (SV40 IMhu), and astrocytes (SVG p12) under standardized in vitro conditions. Cells were maintained in Minimum Essential Medium (MEM, GIBCO) supplemented with 10% fetal calf serum and 1% penicillin-streptomycin. Cultures were incubated at 37°C with 5% CO_2_. Glioma cells were seeded at a density of 5×10^6^ cells per 75-cm^2^ flask and cultured for 48 hours under standard conditions and subsequently co-cultured with glial cells at a 1:1 ratio, with matched monocultures as controls. Cultures were maintained under hypoxic (1% O_2_) or normoxic conditions for five days, with controlled media replacement, followed by cell harvesting and -80°C storage. See details in Supplementary Methods.

### RNA extraction and bulk RNA-seq library preparation

See details in Supplementary Methods.

### Bulk RNA-seq data analysis

Raw RNA-seq reads underwent quality control (FastQC^22^), trimming (Trimmomatic^23^), alignment to GRCh38 using STAR^24^, followed by TMM normalization and transformation to log_2_CPM for downstream analyses. See details in Supplementary Methods.

### Machine learning-based identification of candidate marker genes

See details in Supplementary Methods.

### Cell culture validation model and maintenance

See details in Supplementary Methods.

### RNA isolation, cDNA synthesis and quantitative real-time PCR

See details in Supplementary Methods.

### Spatial transcriptomics analysis

Space Ranger outputs were processed in ‘Seurat’ with integration using 3000 features, followed by normalization, scaling, and clustering^25^. Spatial hypoxia scores were computed to define regions as true hypoxia (TH; score > μ+1σ), mild hypoxia (MH; μ ≤ score ≤ μ+1σ), and normoxia (NX; score < μ), followed by downstream spatial enrichment analyses. See details in Supplementary Methods.

### Spatially functional neighborhood analysis

See details in Supplementary Methods.

### Cell-cell interaction and deconvolution analysis in stRNA-seq data

We applied ‘COMMOT’^26^ to construct CCC networks based on the custom defined L-R pairs. Cell-type composition was inferred using DOT^27^. See details in Supplementary Methods.

### NMF-based stratification of hypoxia-associated clusters in TCGA cohort

Hypoxia-associated genes were curated and prognostic genes were identified through Cox regression. ‘NMF’^28^ was applied to characterize hypoxia-defined clusters, which were validated by sample-sample Pearson correlation and single sample gene set enrichment analysis (ssGSEA)^29^. See details in Supplementary Methods.

### Multivariate assessment of hypoxia-driven survival outcomes

See details in Supplementary Methods.

### Characterizing the tumor microenvironment in TCGA cohort

Patients were stratified into four clinical subgroups. Differential expression^30^, Gene set enrichment analysis (GSEA)^31^, ESTIMATE-based microenvironment inference^32^, and immune checkpoint profiling^33–36^ were conducted to compare hypoxia-defined groups. See details in Supplementary Methods.

### Decoupling cellular state plasticity in TCGA cohort

GSEA and ssGSEA were performed using QAD-stage signatures, PROGENy model, and NMF-derived MPs. See details in Supplementary Methods.

### Somatic mutation, copy number alteration, and DNA methylation analyses in TCGA cohort

See details in Supplementary Methods.

### Immunofluorescence staining and image processing

See details in Supplementary Methods.

### Statistical analysis

Analyses were performed in RStudio using the 64-bit R architecture (version 4.4.2) and Jupyter Notebook using Python (version 3.10.0). Data normality was assessed using the Shapiro-Wilk test. Parametric comparisons between groups were performed using Student’s t-test, and associations between continuous variables were evaluated using Pearson correlation. Statistical significance was defined as p < 0.05.

## Results

### Hypoxia is associated with an AC-like state poised for EGFR-mediated activation

Investigating cellular hypoxia-response at single-cell resolution is imperative for unraveling its intricate role in tumor microenvironment dynamics and cellular heterogeneity within IDH-mutant gliomas. To achieve this, we integrated scRNA-seq datasets from Wang et al.,^6^ and Abdelfattah et al.,^7^, representing patients diagnosed with IDH-mutant gliomas (n=7) **(Supplementary Table 1, Supplementary Fig. S1A)**. The tumor microenvironment harbored 7 distinct cell types (12,044 cells), with glioma (52.39%) and myeloid (34.57%) populations predominating **(Fig. 1A)**. Focusing on malignant cells, sub-clustering resolved 12 distinct clusters across patients and pathological subtypes **(Supplementary Fig. S1B)**. Unbiased non-negative matrix factorization (NMF) identified shared tumor-associated transcriptional programs. Hierarchical clustering of these programs across patients resolved into four metaprograms: AC-like (*APOE, VIM, ID4*), OPC-like (*CNP, GPR17, ZEB2*), NPC-like (*SOX11, EIF4B, CDH7*), and Cycling (*SLBP, PCNA, HMGB2*) states **(Fig. 1B; Supplementary Fig. S1C-D, Supplementary Table 2)**. These MPs showed significant overlap with canonical glioma states defined by Tirosh et al.,^4^ **(Supplementary Fig. S1C)**. Meta-module scoring of MPs and hypoxia signature showed that NPC-like, OPC-like, and Cycling programs were enriched in distinct non-hypoxic clusters, whereas the AC-like program strongly co-localized with hypoxia-enriched clusters (clusters 1, 7, 4, 11, 0, 8) **(Fig. 1C-D, Supplementary Fig. 1E)**. Consistently, hypoxia response positively correlated with the AC-like program **(Fig. 1E)**. Cells were annotated by dominant meta-module scores as AC-like, OPC-like, NPC-like, and Cycling states. Pathway activity inference revealed a depletion of proliferation-associated pathways (MAPK, PI3K) in the AC-like state, consistent with its less-proliferative nature^3,37,38^. However, elevated JAK-STAT and EGFR signaling suggest this population is transcriptionally poised for activation^39,40^ **(Fig. 1F)**. This upregulation of EGFR signaling was driven by leading-edge genes (*KLHL24, S100A16, SLC15A2, EGR1, EGR2*) **(Supplementary Fig. 1F, Supplementary Table 3)**. Furthermore, high TRAIL, p53, and TGF-β activity in AC-like state indicated a stress-adapted and apoptosis-resistant state^41–43^. Additional enrichment of oxidative phosphorylation, electron transport, ribosomal translation, interferon, and TNF-α/NF-κB signaling indicated a metabolically active and immune-associated properties **(Fig. 1G)**.

**Fig. 1.**
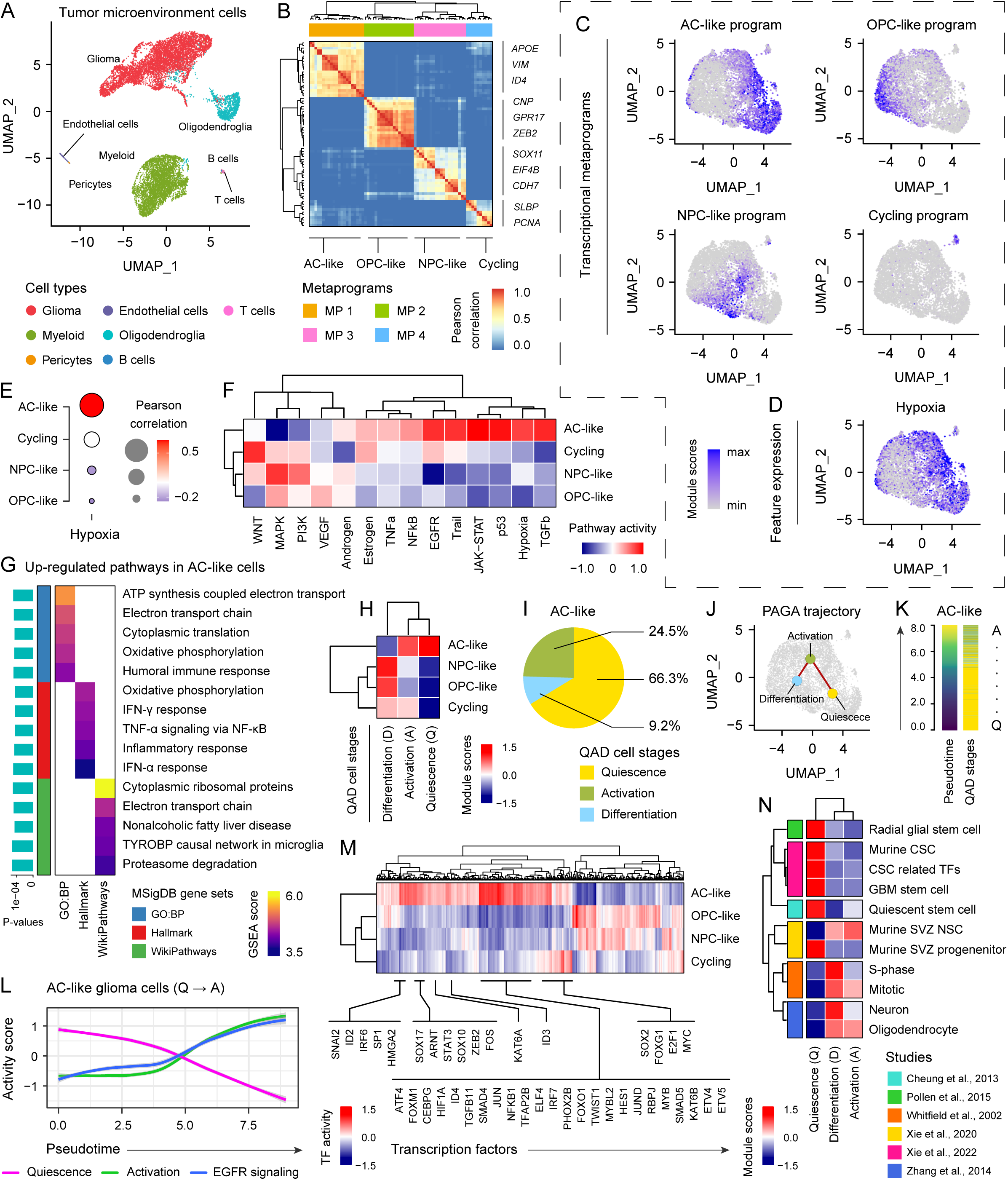
Single-cell profiling reveals hypoxia-associated AC-like cells with quiescent GSC-like features. (A) Dimensional reduction plot (UMAP) of 12,044 single cells depicts the cellular composition of the IDH-mutant glioma microenvironment, where each cell is represented by individual dots and colored according to its cell type. (B) Heatmap displaying the Pearson correlation between gene expression programs derived from sample-wise non-negative matrix factorization (NMF), clustered using Ward’s D2 method to identify four conserved cellular states or metaprograms (MPs). (C-D) UMAP projection of glioma cells highlighting enrichment for four transcriptional metaprograms (AC-like, OPC-like, NPC-like, and Cycling) and hypoxia. Cells are colored based on module enrichment scores for each signature. (E) Bubble plot illustrating the single-cell correlation between hypoxia and the four transcriptional metaprograms. Circle color and size represent the magnitude of the Pearson correlation coefficient (r). (F) Inference of PROGENy pathway activity across cell types assigned to specific transcriptional metaprograms. The heatmap is colored by median enrichment scores. (G) Gene set enrichment analysis (GSEA) for GO:BP, Hallmark, and Reactome signatures, showing the top significantly enriched terms in AC-like glioma cells. Gene counts indicate the number of genes contributing to each enriched pathway or signature. (H) Heatmap depicting enrichment scores for QAD (Quiescence, Activation, Differentiation) stages across annotated glioma MPs. (I) Pie chart representing the percentage of Q-, A-, and D-stages within AC-like glioma cells. (J) Partition-based graph abstraction (PAGA) trajectory projected onto a UMAP embedding of malignant cells. The nodes and connecting edges illustrate the transcriptional transitions and connectivity between QAD stages. (K) Pseudotime ordering of AC-like cells with the corresponding distribution of QAD-stages. (L) Line plots depicting the dynamic changes in EGFR signaling, quiescence, and activation program activities along the pseudotime trajectory in the AC-like glioma population. (M) Heatmap depicting the highest-ranked transcription factors enriched across glioma cell categories. (N) Heatmap showing the enrichment of published gene signatures characterizing the QAD stages in AC-like gliomas.

Next, we characterized the tumors using glioma-associated QAD (GBM QAD) signatures, which include the quiescence (Q-stage), activation (A-stage), and differentiation (D-stage) stages outlined by Foerster et al^5^. The AC-like cells showed elevated expression of Q- and A-stages **(Fig. 1H, Supplementary Fig. 2A-B)**. We also assessed the distribution of QAD stages (annotated based on meta-module score higher than median) across glioma cell categories **(Supplementary Fig. 2C)**, revealing that 66.3% of the AC-like cells remained in a quiescent stage, whereas 24.5% reflected an activated stage **(Fig. 1I, Supplementary Fig. 2D)**, highlighting AC-like gliomas as Q-stage and A-stage dominated tumors. Trajectory inference delineated a continuum spanning these stages, with Q-stage at the root and A-stage downstream **(Fig. 1J; Supplementary Fig. 2E)**. This Q-to-A transition was largely confined to AC-like cells, reflecting high transcriptional plasticity **(Fig. 1K)**. Notably, EGFR signaling activity increased along this trajectory with progression of the activation program, while quiescence declined **(Fig. 1L)**, suggesting its role in initiating the Q-to-A transition within AC-like gliomas and priming cells for proliferation^5^. Transcription factor inference revealed enrichment of hypoxia (HIF1A, ATF4), quiescence/GSC (SOX2, SNAI2), proliferation (E2F1, MYC), and NOTCH signaling-related regulators (ID2, ID3) in AC-like gliomas **(Fig. 1M)**. Additionally, Q-stage strongly overlapped with established glioma stem cells (GSC) and neural stem cell signatures^13–17^, whereas A-stage associated with proliferating progenitors^17^ **(Fig. 1N)**. Collectively, these findings support the characterization of a hypoxia-associated AC-like state as a quiescent, GSC-like reservoir poised for EGFR-mediated activation and potential progression toward proliferative states.

### Microglia-derived HBEGF activates EGFR signaling in hypoxic AC-like gliomas

Intercellular communication within the glioma microenvironment plays a critical role in modulating tumor cell behavior and progression. Given that elevated EGFR signaling is implicated in initiating the Q-to-A transition in AC-like gliomas, we sought to identify microenvironmental cues influencing this process. We first re-clustered the myeloid population from the same studies (Wang et al.,^6^ and Abdelfattah et al.,^7^) **(Supplementary Fig. 3A)** and classified them into four categories: microglia (81.14%), macrophages (14.73%), myeloid-derived suppressor cells (MDSCs) (2.74%), and dendritic cells (DCs) (1.38%) **(Supplementary Fig. 3B-C)**. Using CellChat^20^, we deciphered interactions between immune (myeloid and lymphoid populations) and malignant cells **(Supplementary Fig. 3D)**. Global signaling activity identified microglia, AC-like, and OPC-like glioma cells as the dominant signaling partners **(Fig. 2A)**. To delineate immune-to-glioma influence, we quantified paracrine communication directionality. Among all immune cell types, microglia displayed relatively greater interaction number and strength toward the AC-like population, underscoring their dominant role in tumor-immune crosstalk **(Fig. 2B-C)**. We identified 18 distinct ligand-receptor (L-R) pathways mediating immune-glioma interactions, among which microglia prominently engaged in SPP1, PDGF, IGF, EGF, and CypA signaling **(Fig. 2D)**. Crucially, microglia functioned as the principal sender and influencer of the EGF pathway, while AC-like cells acted as the primary receivers with the highest communication probabilities **(Fig. 2E, Supplementary Fig. 3E)**. In contrast, SPP1, PDGF, IGF, and CypA pathways showed weaker microglial contribution, with other cell types driving these interactions **(Supplementary Fig. 3F)**.

**Fig. 2.**
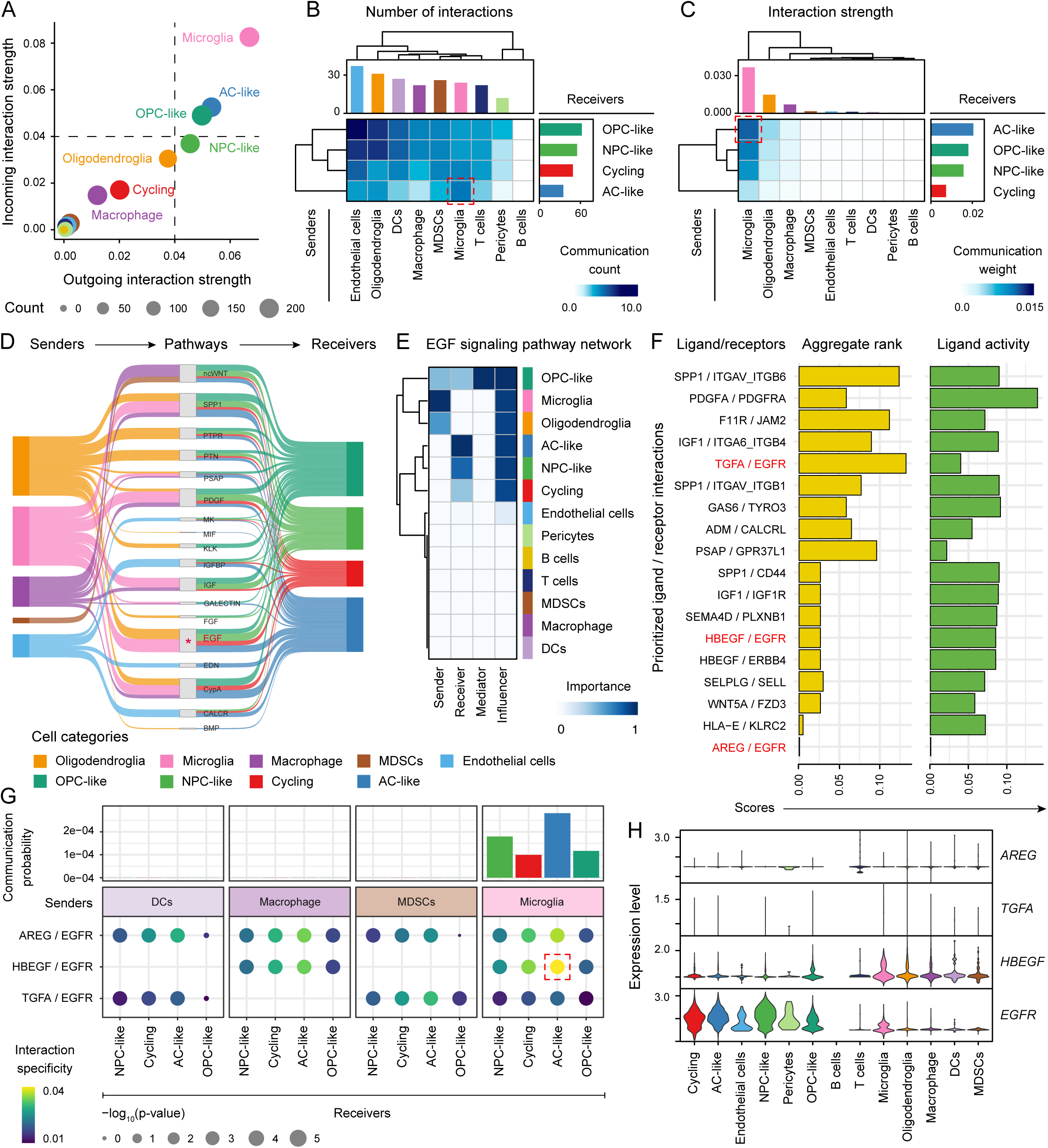
Cell–cell communication reveals microglia–glioma crosstalk. (A) 2D scatter plot depicting the intensity of outgoing or incoming interactions of glioma and non-malignant cell types. Dot size is proportional to the number of significantly expressed receptor-ligand pathways in different cell populations. (B-C) Heatmaps showing interaction counts and strengths of secretory pathways mediating signaling from the non-malignant populations (senders) to glioma cell populations (receivers). The top and right-colored bar plots indicate the total number of interactions for each cell group. (D) Inferred intercellular signaling pathways from immune cells (senders) to glioma cell populations (receivers). The thickness of each flow represents the relative contribution of each cell group to the signaling pathways. (E) Heatmap showing the relative importance of each cell group as senders, receivers, mediators, and influencers within the EGF signaling network. (F) Top ligand-receptor pairs mediating communication from microglia (senders) to AC-like glioma cells (receivers). Ligand-receptor pairs highlighted in red are associated with the EGF signaling pathway. (G) Dot plot illustrating EGF pathway-specific ligand-receptor pairs mediating signaling from myeloid populations (senders) to glioma subtypes (receivers). Dot color indicates interaction specificity, and dot size represents −log_10_(p-value) (statistical significance). The upper bar plot displays the communication probability of each ligand-receptor pair. (H) Violin plots illustrating the expression distribution of ligand-receptor genes contributing to the inferred EGF signaling network across cell populations.

We next aimed to decipher the potential L-R interactions between microglia and the AC-like glioma population. Among top interactions, SPP1/ITGAV-ITGB6 and PDGFA/PDGFRA were identified, while multiple EGF ligands (HBEGF, AREG, TGFA) engaged EGFR, with the HBEGF/EGFR pair displaying strong and significant interaction, characterized by high ligand activity **(Fig. 2F)**. Comparative analysis of EGF ligands confirmed that the HBEGF/EGFR interaction displayed the highest communication and interaction specificity when microglia-derived HBEGF interacted with EGFR expressed by AC-like glioma cells **(Fig. 2G)**. Gene expression analyses further confirmed elevated expression of HBEGF in microglia and EGFR in AC-like glioma cells, reinforcing the likelihood of a microglia-driven HBEGF/EGFR signaling axis **(Fig. 2H)**.

Independent validation using ICELLNET^44^, with immune cells defined as senders (surrounding condition) and AC-like gliomas as receivers (central condition) under an inward communication framework **(Supplementary Fig. 4A)**, confirmed predominant EGFR-family ligand signaling from microglia **(Supplementary Fig. 4B)**, with the HBEGF/EGF pair exhibiting the strongest communication with AC-like cells **(Supplementary Fig. 4C)**. Collectively, these results identify microglia as key modulators of the glioma microenvironment, providing HBEGF-mediated paracrine signals that activate EGFR signaling and potentially prime quiescent AC-like glioma cells for activation.

### Microglia promote an AC-like program under hypoxia

Previous studies have highlighted the roles of microglia in shaping the glioma TME^45,46^. To investigate their contributions and experimentally contextualize our findings, we established a primary IDH-mutant glioma cell line (UKFR #971) retaining the IDH-R132H mutation **(Supplementary Fig. 5A)**. Glioma cells were cultured under monoculture (baseline control) and co-culture conditions with microglia (positive control) or astrocytes (negative control), under normoxic and hypoxic environments. Differential expression analysis comparing hypoxic and normoxic conditions across tumor monoculture (Tum^H+^ vs Tum^H-^), microglia/tumor co-culture (Tum/Mic^H+^ vs Tum/Mic^H-^), and astrocyte/tumor co-culture (Tum/Ast^H+^ vs Tum/Ast^H-^) was performed **(Fig. 3A)**. In monoculture, 662 genes were upregulated under hypoxia (Tum^H+^), including ***CHRDL1*, *SFRP2*, and *CXCL12* (Fig. 3B)**. In both co-culture conditions under hypoxia Tum/Mic^H+^ and Tum/Ast^H+^), significantly elevated genes included *VEGFA*, *IGFBP3*, and *IGFBP5* **(Fig. 3C-D)**. GSEA revealed that hallmark pathways, including hypoxia, EMT, angiogenesis, IL2-STAT5, and TNF-α/NF-κB signaling, were strongly upregulated in the Tum/Mic^H+^ condition, whereas Tum/Ast^H+^ showed only modest enrichment of hypoxia and myogenesis pathways **(Fig. 3E)**. To evaluate the effects of microglia and astrocytes on glioma, independent of hypoxia, we compared Tum^H+^ with Tum/Mic^H+^ and Tum/Ast^H+^ conditions **(Fig. 3A, Supplementary Fig. 5B-C)**. Notably, cell cycle-associated pathways were significantly upregulated in Tum/Mic^H+^ condition, whereas no hallmark pathways were significant in Tum/Ast^H+^ condition **(Supplementary Fig. 5D)**, highlighting the dominant modulatory role of microglia in IDH-mutant gliomas.

**Fig. 3.**
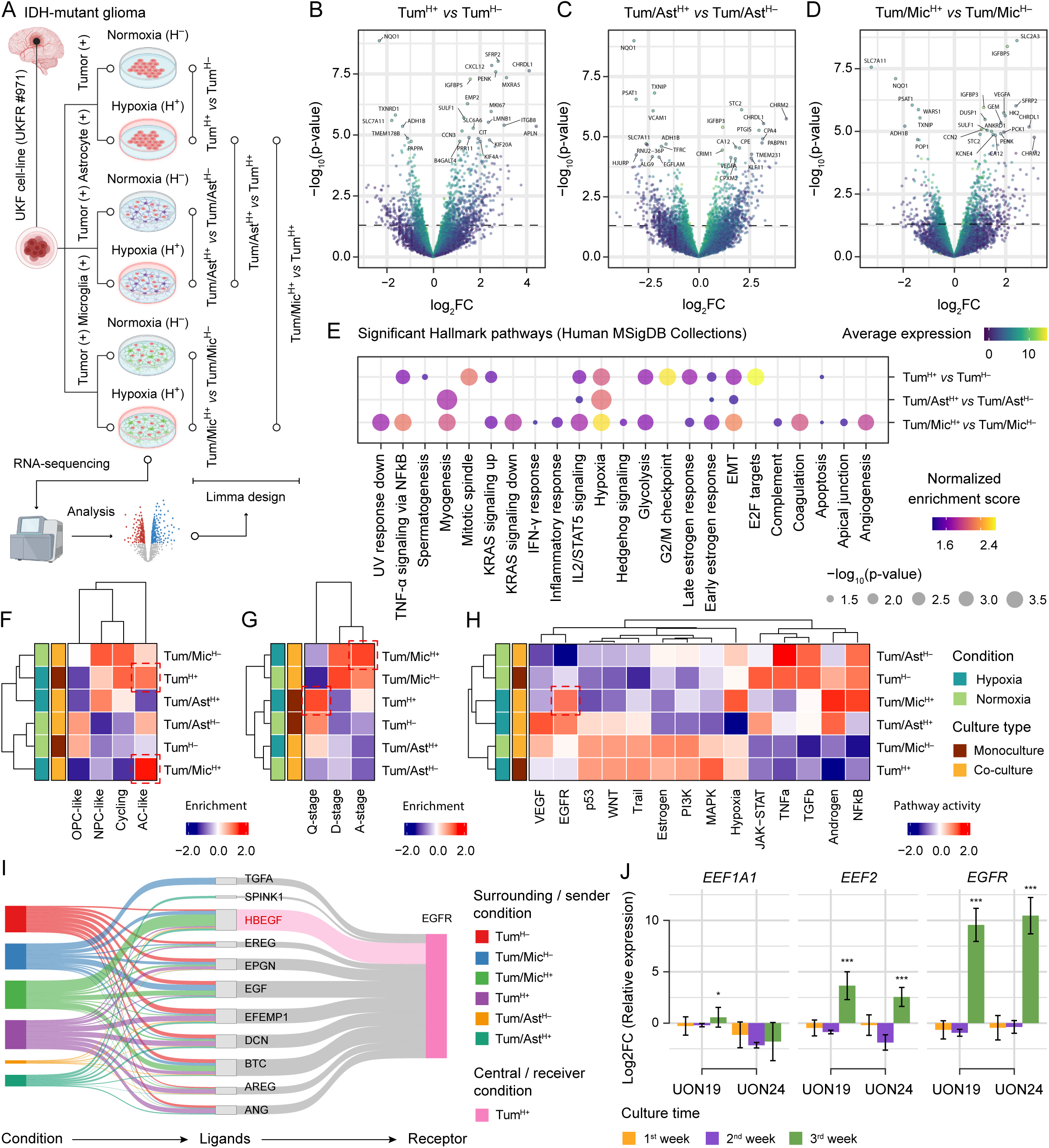
Co-culture models uncover microenvironmental factors shaping the cellular programs in IDH-mutant glioma. (A) Schematic representation of monoculture and co-culture experimental setup, followed by bulk RNA sequencing and transcriptomic analysis. Created with BioRender.com. (B-D) Volcano plots depicting differential gene expression between hypoxic and normoxic conditions in tumor monoculture (Tum^H+^ vs Tum^H-^), astrocyte/tumor co-culture (Tum/Ast^H+^ vs Tum/Ast^H-^), and microglia/tumor co-culture (Tum/Mic^H+^ vs Tum/Mic^H-^), respectively. (E) Gene set enrichment analysis (GSEA) of significantly enriched Hallmark pathways (p<0.05) across the three differential expression conditions. Dot color represents the normalized enrichment score (NES), and dot size indicates −log_10_(p-value) (statistical significance). (F-G) Heatmap showing median enrichment scores for IDH-mutant glioma MPs and QAD (stages across each culture condition. (H) Heatmap showing the differences in pathway activity inference for PROGENy biological pathways between each culture condition. (I) Sankey diagram illustrating the contributing ligand-receptor interactions and their communication scores from all conditions (sender conditions) toward the hypoxic tumor monoculture condition (receiver condition). The width of each flow is proportional to the communication score, representing the strength of the inferred interaction. (J) Bar plots showing RT-qPCR-derived log_2_ fold change (log_2_FC) values for *EEF1A1*, *EEF2*, and *EGFR* expression in UON19 and UON24 cell lines under hypoxic conditions at weeks 1, 2, and 3. Expression levels are presented as log_2_FC relative to normoxic controls. Error bars represent the standard error of the mean of biological replicates (n=9). P-values are represented by asterisks: *** (p < 0.001), ** (p < 0.01), and * (p < 0.05).

Next, we investigated the influence of microglia and astrocytes on glioma MPs and QAD stages in both hypoxic and normoxic conditions. GSEA demonstrated that the AC-like program was enriched in Tum^H+^ and Tum/Mic^H+^ conditions **(Fig. 3F)**, indicating its dependence on hypoxia and modulation by microglial interactions. To assess whether Q-to-A dynamics followed a similar pattern, we performed enrichment analysis using a similar approach. The Q-stage was upregulated in Tum^H+^, whereas the A-stage was strongly enriched in the Tum/Mic^H+^ condition **(Fig. 3G)**, supporting our earlier observation that hypoxia primarily maintains a quiescent AC-like state, while microglia promote the transition toward an activated phenotype. Consistently, pathway activity inference confirmed selective elevation of EGFR signaling in the Tum/Mic^H+^ condition **(Fig. 3H)**. Cell-cell interaction analysis using ICELLNET^44^, with Tum^H+^ as receiver (central condition) and all culture conditions as senders (surrounding condition), with an inward communication direction **(Supplementary Fig. 5E)**, identified HBEGF as a potential ligand in the Tum/Mic^H+^ condition interacting with EGFR on glioma cells in Tum^H+^ condition **(Fig. 3I)**, supporting a microglia-driven paracrine HBEGF/EGFR signaling.

To prioritize MP-associated markers representing both Tum^H+^ and Tum/Mic^H+^ conditions, differential expression was integrated with machine learning predictions, identifying EEF2 as an AC-like marker **(Supplementary Fig. 5F)**. To validate our findings, primary IDH-mutant glioma cell lines (UON19 and UON24) were cultured, and gene expression levels of *EEF2* (a hypoxia-responsive AC-like marker), *EGFR*, and *EEF1A1* (a translational marker preferentially expressed in NPC-like cells) were quantified by qPCR at 1-, 2-, and 3-week time points under normoxic and hypoxic conditions **(Supplementary Fig. 5G)**. After an initial acute stress response, both *EEF2* and *EGFR* were highly up-regulated by the third week of hypoxia, coinciding with cellular adaptation **(Fig. 3J)**. Collectively, these findings demonstrate that hypoxia maintains IDH-mutant gliomas in a quiescent AC-like state, while microglia-derived HBEGF/EGFR signaling promotes their transition to activation, highlighting a paracrine mechanism.

### AC-like gliomas selectively localize and activate within hypoxic niches

To investigate the spatial localization of hypoxic regions and their association with the MPs, we leveraged spatial transcriptomics (stRNA-seq) datasets^8,9^ from eight IDH-mutant glioma patients (n=8) **(Supplementary Table 4)**. We identified 4-11 Seurat clusters across samples **(Supplementary Fig. 6A)**. Integrated analysis classified 21,261 spots into true hypoxia (TH), mild hypoxia (MH), and normoxia (NX) based on enrichment scores **(Fig. 4A-B)**. Of the total 21,261 barcode spots, 2,682 (12.61%) exceeded one standard deviation above the median enrichment score and were classified as true-hypoxia; 7,949 (37.39%) spots between the median and one standard deviation above the median were categorized as mild-hypoxia; and 10,630 (49.99%) spots below the median value were classified as normoxia **(Fig. 4B)**. Five samples (BWH23, BWH24, BWH25, LG1, LG2) exhibited higher proportions of TH regions **(Supplementary Fig. 6B)**, with BWH24, BWH23, and BWH25 showing highest percentage of true-hypoxia spots with centered distribution **(Fig. 4C-D, Supplementary Fig. 6C)**. Other samples displayed diffused hypoxia **(Supplementary Fig. 6D)**, indicating centered hypoxic regions are not universal in IDH-mutant gliomas.

**Fig. 4.**
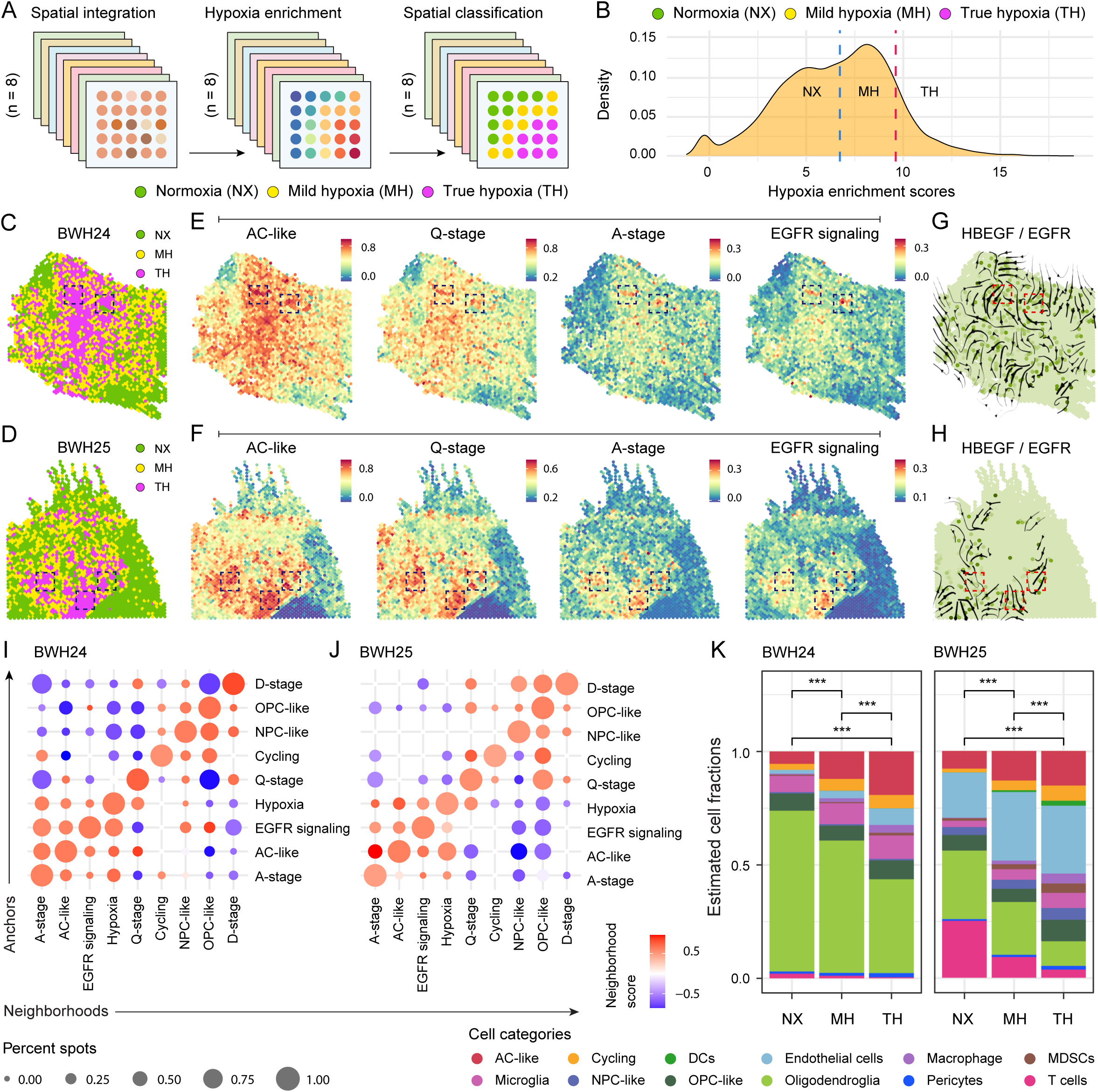
Spatial transcriptomics reveals the localization of cellular programs within hypoxic niches. (A) Schematic representation of stRNA-seq dataset integration, enrichment, and classification of barcode spots into three hypoxic regions (true hypoxia, mild hypoxia, and normoxia). (B) Density plot showing the distribution of hypoxia enrichment across all spatial samples. The blue dashed line marks the median hypoxia enrichment level, while the red dashed line represents one standard deviation above the median. (C-D) Spatial feature plot showing the distribution of classified hypoxic regions in samples BWH24 and BWH25, with a centralized arrangement of the true hypoxic (TH) region. (E-F) Spatial GSEA surface plots illustrating enrichment scores for the AC-like program, quiescence (Q-stage), activation (A-stage), and EGFR signaling in samples BWH24 and BWH25, respectively. Each spatial spot is colored according to its enrichment score, where dark red denotes regions with strong enrichment and dark blue indicates regions with minimal or no enrichment. (G-H) Directionality and intensity of HBEGF/EGFR signaling in samples BWH24 and BWH25, respectively. Arrow direction indicates the source-to-target signaling flow, and arrow thickness represents the relative intensity of the inferred signal. (I-J) Dot plots illustrating the spatial co-localization of the transcriptional features (hypoxia, AC-like, OPC-like, NPC-like, Cycling, QAD-stages, and EGFR signaling) in the neighboring spots for samples BWH24 and BWH25, respectively. Dot color represents the co-occurrence values between pairs of transcriptional features, while dot size indicates the percentage of co-occurrence within the same spatial neighborhood. (K) Stacked bar plots showing the relative abundances of cell types within decomposed spatial spots classified as hypoxic regions in samples BWH24 and BWH25, respectively. Cell type proportions were inferred based on reference scRNA-seq data.

GSEA revealed strong enrichment of the AC-like program within TH regions in BWH23, BWH24, and BWH25 **(Fig. 4E-F, Supplementary Fig. 7A)**. In BWH24 and BWH25, both the AC-like program and Q-stage were highly enriched in centralized TH regions, whereas other signatures were less prominent **(Fig. 4E-F, Supplementary Fig. 7B-C)**. Within these TH regions, we identified discrete A-stage niches marked by pronounced EGFR signaling, occurring concurrently with AC-like and Q-stage **(Fig. 4E-F)**. In contrast, Q-stage regions were largely located outside TH zones in BWH23, reflecting patient-specific heterogeneity **(Supplementary Fig. 7A)**. Samples with diffuse hypoxia showed minimal AC-like enrichment and lacked EGFR signaling activity **(Supplementary Fig. 7D-H)**. Inference of HBEGF/EGFR signaling directionality in BWH24 **(Fig. 4G)** and BWH25 **(Fig. 4H)** revealed activated spots within the niche that predominantly received signals from neighboring areas, suggesting these sites function as seeding sites for the Q-to-A transition.

To quantitatively measure the overlapping neighborhoods, we implemented a Spatially Functional Neighborhood Analysis where the anchors are the gene signatures for which enriched spots (selected from the distribution of the enrichment score) were used to define the neighborhood spanning across a 250µm boundary. The neighborhood gene signatures are those for which the enrichment within the anchored neighborhood is calculated. In BWH24 and BWH25, the hypoxia-anchored neighborhood exhibited a 50% overlap with AC-like and Q-stage spots, consistent with prior findings **(Fig. 4I-J)**. It indicated these molecular features are spatially associated, potentially reflecting a functional relationship. Furthermore, 15-25% of AC-like and A-stage spots overlapped with EGFR signaling neighborhoods **(Fig. 4I-J)**, confirming localized activation within these niches. In samples with lower TH proportions, these molecular features were largely non-overlapping **(Supplementary Fig. 8A)**.

To uncover the cellular composition of the spatially classified regions, we performed single-cell deconvolution using datasets from Wang et al.,^6^ and Abdelfattah et al.,^7^. In BWH24 and BWH25, we observed a significant progressive increase in AC-like glioma cells from NX to MH, with greater abundance in TH regions. Notably, microglia were also enriched in TH regions **(Fig. 4K)**, supporting their role in the Q-to-A transition. In contrast, BWH23 showed a uniform distribution of AC-like cells **(Supplementary Fig. 8B)**. Together, these findings establish that hypoxic niches serve as reservoirs for the AC-like state, where microglia-derived HBEGF/EGFR signaling drives quiescent cells toward activation, representing critical sites for tumor persistence and progression.

### Hypoxia is elevated in higher-grade gliomas and sustains the AC-like state

To investigate the impact of hypoxia across a larger cohort, we analyzed 415 IDH-mutant cases, comprising both 1p/19q-codeletion (Oligodendroglioma) and 1p/19q-non-codeletion (Astrocytoma), from the ‘Brain Lower Grade Glioma (TCGA, PanCancer Atlas)’ cohort. Astrocytomas of WHO grade 4 were excluded from the study. To identify clinically relevant hypoxia-associated genes, the MSigDB hallmark hypoxia set was assessed and 35 genes significantly associated with poor prognosis were selected **(Supplementary Table 5)**. The expression profile of these genes (n=35) was then subjected to NMF clustering, which optimally resolved two molecular subtypes (k=2) based on cophenetic, dispersion, and silhouette metrics **(Supplementary Fig. 9A-B)**. Samples were stratified into C1 (n=293) and C2 (n=122), with strong intra-cluster coherence **(Fig. 5A)**. ssGSEA showed that the hypoxia gene signature was enriched in the C2 group, hence termed as “severe hypoxia”, compared to the C1 group termed as “mild hypoxia” **(Fig. 5B)**. Severe hypoxic patients were associated with significantly reduced overall survival (OS) (log-rank p=0.0011) **(Fig. 5C)**.

**Fig. 5.**
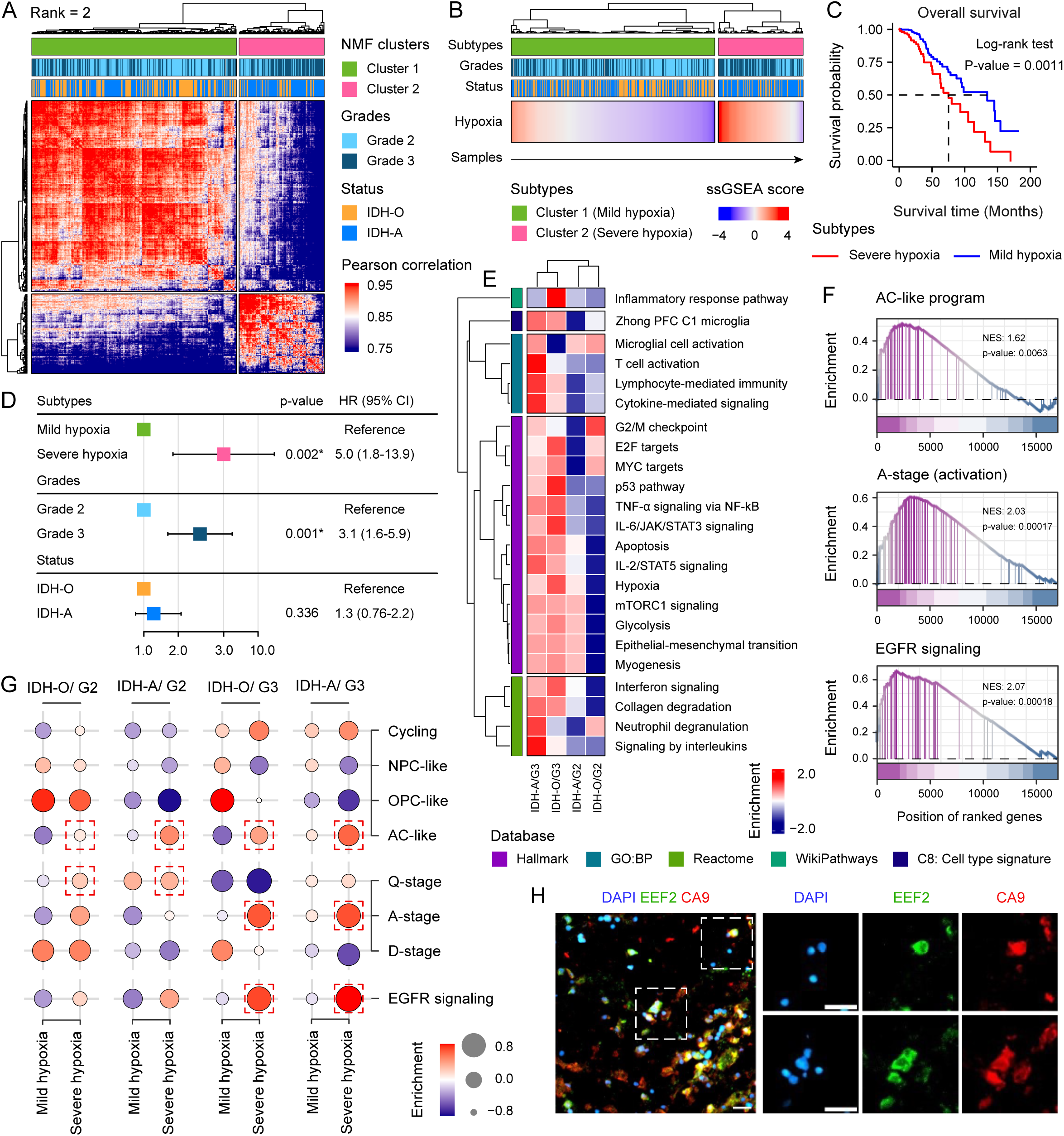
Large-scale characterization of hypoxia-associated transcriptional programs across tumor grades and molecular subtypes. (A) Heatmap representing the sample-sample correlation matrix, stratified according to non-negative matrix factorization (NMF) at rank 2 (k=2). Pearson correlation coefficients were computed across the entire cohort to assess the similarity between samples. Both rows and columns are ordered through hierarchical clustering based on the average Euclidean distance. (B) Single-sample gene set enrichment analysis (ssGSEA) for the Hallmark Hypoxia gene set, comprising hypoxia-associated genes significantly correlated with poor prognosis (p<0.05). The heatmap depicts enrichment scores across all samples and is stratified according to NMF-identified molecular subtypes. (C) Kaplan-Meier curve representing overall survival between the two subtypes with a log-rank p-value. (D) Forest plot of hazard ratios from a multivariate Cox proportional hazards model evaluating the prognostic impact of hypoxia status, adjusted for WHO grade and molecular tumor type. (E) Pathway enrichment analysis using Hallmark, GO:BP, Reactome, WikiPathways, and cell type signature gene sets, showing the most significantly enriched biological pathways within the severe hypoxic subtype across IDH-O/G3, IDH-A/G3, IDH-A/G2, and IDH-O/G2 tumor groups. (F) GSEA plots of ranked genes showing enrichment of the AC-like program (top), activation stage (middle), and EGFR signaling (bottom) in severe hypoxia compared to the mild hypoxic group. (G) ssGSEA of IDH-mutant glioma MPs, QAD stages, and EGFR signaling across IDH-O/G2, IDH-A/G2, IDH-O/G3, and IDH-A/G3 tumors stratified by mild and severe hypoxia. Circle color and size represent the magnitude of enrichment. (H) Representative immunofluorescence images of IDH-mutant glioma sample stained for DAPI (nuclei), EEF2 (hypoxia-responsive AC-like marker), and CA9 (hypoxia marker). Scale bars: 25µm.

Clinically, the severe hypoxic group was characterized by a higher prevalence of grade-3 astrocytomas (1p/19q-non-codeletion) **(Supplementary Fig. 10A)**. Multivariable Cox regression analysis revealed that both severe hypoxia and higher tumor grade were associated with poorer overall survival, indicating that tumor grade substantially contributed to the adverse prognosis observed in the hypoxic subgroup **(Fig. 5D)**. Genomic analysis identified an arm-level loss of chromosome 11p in the severe hypoxia subtype, suggesting this alteration may contribute to its distinct molecular profile **(Supplementary Fig. 10B)**. Mutational profiling showed no significant genomic divergence between the hypoxic subtypes, with both groups exhibiting canonical alterations of IDH-mutant gliomas, including *IDH1*, *TP53*, and *ATRX* mutations **(Supplementary Fig. 11A-B)**. The severe hypoxic group exhibited a significantly higher mutational burden (p=3.6×10^-5^) **(Supplementary Fig. 11C)** and lower global methylation levels (p=0.002) **(Supplementary Fig. 11D)**, consistent with its association with higher tumor grade and loss of global methylation^47^. Therefore, these hypoxic subtypes largely retained the fundamental genomic and epigenomic landscape of IDH-mutant gliomas^47^, with no distinctive alterations observed, suggesting that genomic and epigenomic variability minimally contributes to their defining characteristics.

Next, we investigated the role of TME in defining the characteristics of hypoxic subtypes across four clinical categories: oligodendroglioma grade-2 (IDH-O/G2), oligodendroglioma grade-3 (IDH-O/G3), astrocytoma grade-2 (IDH-A/G2), and astrocytoma grade-3 (IDH-A/G3). It revealed a greater fold-change amplitude of significantly upregulated genes in grade-3 tumors under severe hypoxia **(Supplementary Fig. 12A)**. GSEA demonstrated that severe hypoxia in higher-grade gliomas is characterized by microglial activation, cytokine signaling, glycolysis, and oncogenic proliferative programs (E2F and MYC targets, G2/M checkpoint) **(Fig. 5E)**. Furthermore, severe hypoxic gliomas, particularly astrocytomas, showed higher immune and stromal infiltration along with elevated immune checkpoint expression (*CTLA4*, *PDCD1LG2*, *HAVCR2*)^33–36^, that presumably reflected the increased immune cell abundance **(Supplementary Fig. 12B-C, Supplementary Table 6)**.

Next, we uncovered subtype-specific differences in MPs, QAD stages, and EGFR signaling activity. The severe hypoxic group exhibited significant enrichment of A-stage, EGFR signaling, and AC-like and Cycling programs **(Fig. 5F, Supplementary Fig. 12D)**. Tumor type- and grade-specific insights revealed that while the AC-like state was consistent across the severe hypoxia subgroup, the Q-stage was observed in low-grade tumors, whereas the A-stage coupled with EGFR signaling was enriched in higher-grade tumors **(Fig. 5G)**, indicating EGFR-driven Q-to-A transition linked to progression. Finally, immunofluorescence staining in four grade-3 IDH-mutant gliomas demonstrated strong co-localization of EEF2 (hypoxia-responsive AC-like marker) with the tumor-associated hypoxia marker CA9, confirming spatial coupling of hypoxia and AC-like state **(Fig. 5H, Supplementary Fig. 13A-B)**. These findings indicate that hypoxia sustains a quiescent AC-like state in lower-grade tumors, while in higher-grade gliomas, it facilitates an EGFR-driven transition to an activated, proliferative state, driving tumor progression.

## Discussion

Tumor biology is shaped by a complex interplay of genetic, epigenetic, and microenvironmental factors, among which hypoxia is a critical stressor that alters glioma cell behavior and transcriptional states. Investigating IDH-mutant gliomas within the framework of Tirosh et al^4^, we identified a hypoxia-associated AC-like population exhibiting a dual transcriptional profile, comprising Q- and A-stages. The overlap of Q-stage with GSC-like signatures suggests these cells harbor features of a quiescent reservoir, often refractory to conventional treatments^16^. Conversely, A-stage displayed strong enrichment of proliferative neural progenitor signatures **(Fig. 1)**. This bifurcation highlights a fundamental aspect of transcriptional plasticity where dormant, GSC-like cells are primed for activation, consistent with studies on dynamic transitions regulated by NOTCH and WNT pathways promoting tumor heterogeneity and therapeutic resistance^48^. While Foerster et al.,^5^ characterized the Q-to-A transition to aberrant WNT signaling underlying GBM progression, our study extends these findings by incorporating microenvironmental stressors, such as hypoxia. We demonstrate that while WNT signaling is enriched in the Cycling population, the AC-like population is co-enriched with hypoxia and EGFR signaling **(Fig. 1)**. Within the hypoxic microenvironment, the Q-to-A transition is coupled with EGFR signaling induction and the downregulation of quiescence programs, suggesting EGFR upregulation as a priming mechanism for cellular activation, enabling cells to exit dormancy while selectively retaining quiescence-associated stemness.

Importantly, within established transcriptional programs proposed by Nomura et al.^49^ and Wu et al.^50^, the glial progenitor cell-like (GPC-like) state is defined as a proliferative intermediate positioned between OPC-like/NPC-like programs and the AC-like state. However, we did not identify a transcriptionally distinct GPC-like population in our IDH-mutant glioma cohort. One possibility is that such a program exists but was underrepresented in our dataset. Alternatively, progenitor-associated activation features may be partially embedded within the AC-like A-stage population **(Fig. 1)**, thereby diminishing the emergence of a clearly segregated GPC-like transcriptional state. This interpretation is consistent with recent epigenomic findings by Wu et al.^50^, suggesting that malignant progenitors in IDH-mutant gliomas share a permissive chromatin landscape that enables rapid transitions between developmental programs. It also demonstrated that IDH-mutant glioma cells lack the clear chromatin specification observed in normal neural lineages, allowing tumor cells to dynamically cycle between developmental phenotypes. Within this framework, the hypoxia-associated A-stage may represent a microenvironmentally induced expansion toward such proliferative progenitor-like states within the AC-like compartment.

EGFR signaling networks play a pivotal role in glioma pathogenesis and progression through multiple mechanisms, including receptor overexpression or amplification^51^. While EGFR mutation and amplification are hallmarks of higher-grade GBM^52^, they occur far less frequently in IDH-mutant gliomas^53^. Consistent with these observations, analysis of mutational and CNA from the ‘Brain Lower Grade Glioma (TCGA, PanCancer Atlas)’ cohort revealed an absence of EGFR mutation or amplification in chromosome 7p in IDH-mutant gliomas **(Supplementary Fig. 10-11).** Critically, these cell-intrinsic genomic profiles fail to account for the robust, context-dependent EGFR signaling signatures we detected under hypoxic conditions, suggesting an alternative, non-genomic mode of activation.

We identified microglia as the primary microenvironmental source of EGFR-activating cues, acting as paracrine signals to induce Q-to-A transitions in AC-like glioma cells **(Fig. 2 and 3)**. Microglia are critical TME components influencing tumor growth and resistance through complex signaling networks^54–56^. Specifically, microglia-derived HBEGF promotes EGFR upregulation in AC-like cells, facilitating activation within hypoxic niches. HBEGF, produced by both tumor cells^51^ and stromal components^49,57^, is consistently elevated in GBM and functions as a critical driver of tumor cell proliferation and therapy resistance^58,59^. Notably, macrophage-secreted HBEGF promotes aggressive MES-like GBM state in mouse models^57^. Moreover, HBEGF/EGFR crosstalk drives quiescent neural stem cells (NSCs) toward a reactive phenotype, further underscoring its role in adverse cellular state transitions^60^. Collectively, microglia-derived HBEGF/EGFR signaling represents a crucial mechanism driving tumor plasticity and aggressiveness.

The current paradigm of IDH-mutant glioma progression proposed by Wu et al.^50^, describes a transition from early-stage, slow-cycling OPC-like tumor cells sustained by DNA hypermethylation, to highly proliferative NPC-like populations in advanced disease. As this hypermethylated landscape erodes during progression, tumors acquire compensatory genetic alterations, including amplifications of *PDGFRA*, *MYCN*, and *CDK4* and deletions of *CDKN2A*, that support the expansion of these proliferative progenitor states. Our analysis of the TCGA cohort recapitulates this pattern among patients with lower hypoxia, as OPC-like programs were more prominent in lower-grade, and NPC-like states were enriched in higher-grade gliomas. However, in patients characterized by severe hypoxic responses, we observed a distinct pattern in which the AC-like state was consistently represented across tumor grades and histopathological subtypes. Within the severe hypoxic group, the Q-stage was detected across low-grade tumors, whereas the A-stage, characterized by elevated EGFR signaling, was preferentially enriched in higher-grade tumors, further supporting the bifurcation of the AC-like program **(Fig. 5)**. Our findings therefore complement the current developmental hierarchy by indicating that a hypoxic microenvironment represents an additional layer shaping cellular transitions during tumor evolution.

Notably, large-cohort analysis confirms that severe hypoxia remains a potent indicator of poor prognosis, even when accounting for tumor grade **(Fig. 5D)**. A key contribution of this study is the identification of the EGFR-driven Q-to-A transition as the cellular mechanism underlying this clinical decline. Our study defines how hypoxia contributes to IDH-mutant glioma biology, not simply as a driver of stress adaptation, but as an orchestrator of transcriptional plasticity. By uncovering this mechanism, our findings open new avenues for innovative differentiation therapies designed to interrupt the cycle of dormancy and reactivation. Targeting these adaptive tumor niches may represent a new therapeutic frontier for IDH-mutant gliomas, with the potential to fundamentally alter disease trajectory and extend patient survival.

## Required statements

## Author contributions

D.D., R.R., K.J., G.A., V.M.R., and S.C. conceptualized the project and designed the experiments. D.D., S.M.A., and N.H. curated the data. I.H., S.J.S., J.G., I.V., R.R. (Roelz R.), J.N., R.R. (Rahman R.), M.B., K.J., V.M.R., and S.C. contributed to the resources and materials. V.J., A.P. conducted the cell-culture, co-culture, bulk RNA-seq, and qPCR experiments. N.S.G. carried out the immunofluorescence experiments. A.O.M. established the UKF cell lines, and M.A.S. performed validation through specific staining. D.D. developed the computational pipeline for analyzing bulk RNA-seq, single-cell RNA-seq (scRNA-seq), spatial transcriptomics (stRNA-seq), whole exome sequencing (WES), copy number alterations (CNAs), and DNA methylation datasets. T.D., C.K., G.A., and S.C. developed the methodology for spatially functional neighborhood analysis. D.D., T.D., S.M.A., N.H., K.J., G.A., and S.C performed the formal analysis. D.D. prepared the figures and wrote the original draft of the manuscript. R.H., I.T., T.A.J., M.S., M.P., S.H.J., R.R. (Rahman R.), M.B., J.B., K.J., G.A., V.M.R., and S.C. reviewed and edited the manuscript. D.D., V.J., A.P., N.S.G., and S.C. prepared the final draft. G.A., V.M.R., and S.C. supervised the study. V.M.R acquired the funding. All authors read and approved the final version of the manuscript.

## Conflict of interest statement

The authors declare no competing interests.

## Funding

V.M.R. lab is supported by the Stiftung Sibylle Assmus Foundation and The Ministry of Food, Rural Areas, and Consumer Protection (MLR), Baden-Württemberg (16 (34) – 8402.43 / 0456 E) and DFG MA 10605/1-1. J.B. is supported by BMBF (DiaQNOS: 1010 0665 01). M.B. is supported by the Deutsche Forschungsgemeinschaft (DFG) – CRC1479 (Project ID 441891347-S1), CRC 1160 (Project ID 256073931-Z02), CRC1453 (Project ID 431984000 – S1), TRR167 (Project ID 259373024-Z01), TRR 353/1 (Project ID 471011418-SP02), TRR417 (Project ID: 540805631-S03), FOR 5476 UcarE (Project ID 493802833-P7) and by German Federal Ministry of Education and Research (BMBF), PM4Onco–FKZ 01ZZ2322A, SATURN3 (01KD2206L). S.C., G.A., T.D., and C.K. are supported through MIRACUM consortia funded by the BMBF as part of the Medical Informatics Initiative (EkoEstMed–FKZ 01ZZ2015). The research was supported by other DFG grants: # 520992132, # 441891347 (OncoEscape), # 390939984 (CIBSS), # 272983813 (TRR179), # 256073931 (IMPATH), and # 518316185 (GMCA).

## Ethics statement

The study was conducted in accordance with the Declaration of Helsinki. All tissue samples were obtained under approved Institutional Review Board protocols and appropriate consent at the institutions of origin.

## Data availability

RNA-seq data generated in this study have been deposited at the Gene Expression Omnibus (GEO) repository of the National Center for Biotechnology Information (NCBI) under primary accession numbers GSE319729. All other data generated in this study are included in the manuscript and the Supplementary Information.

## Code availability

The codes generated in this study have been deposited and are publicly available in GitHub (https://github.com/3DBMandNE-Lab/LGG_Hypoxia).

## Notes

### Competing Interest Statement

The authors have declared no competing interest.

