## Supplementary figures and images for "Spatially confined niches support hypoxia-associated transcriptional plasticity contributing to malignant progression in IDH-mutant gliomas"

### Supplementary Figure 1.pdf

#### Supplementary Figure 1

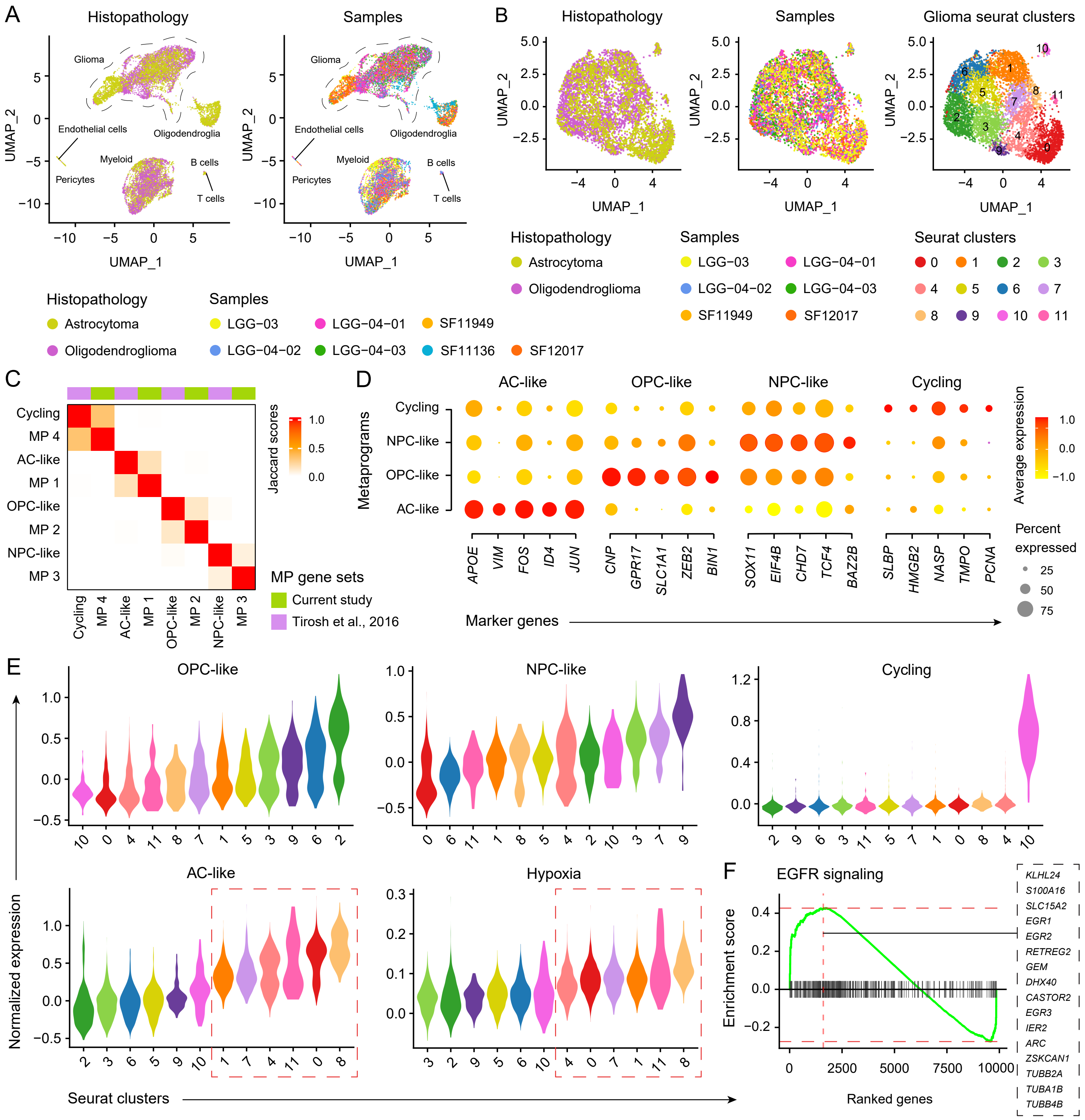

### Supplementary Figure 2.pdf

#### Supplementary Figure 2

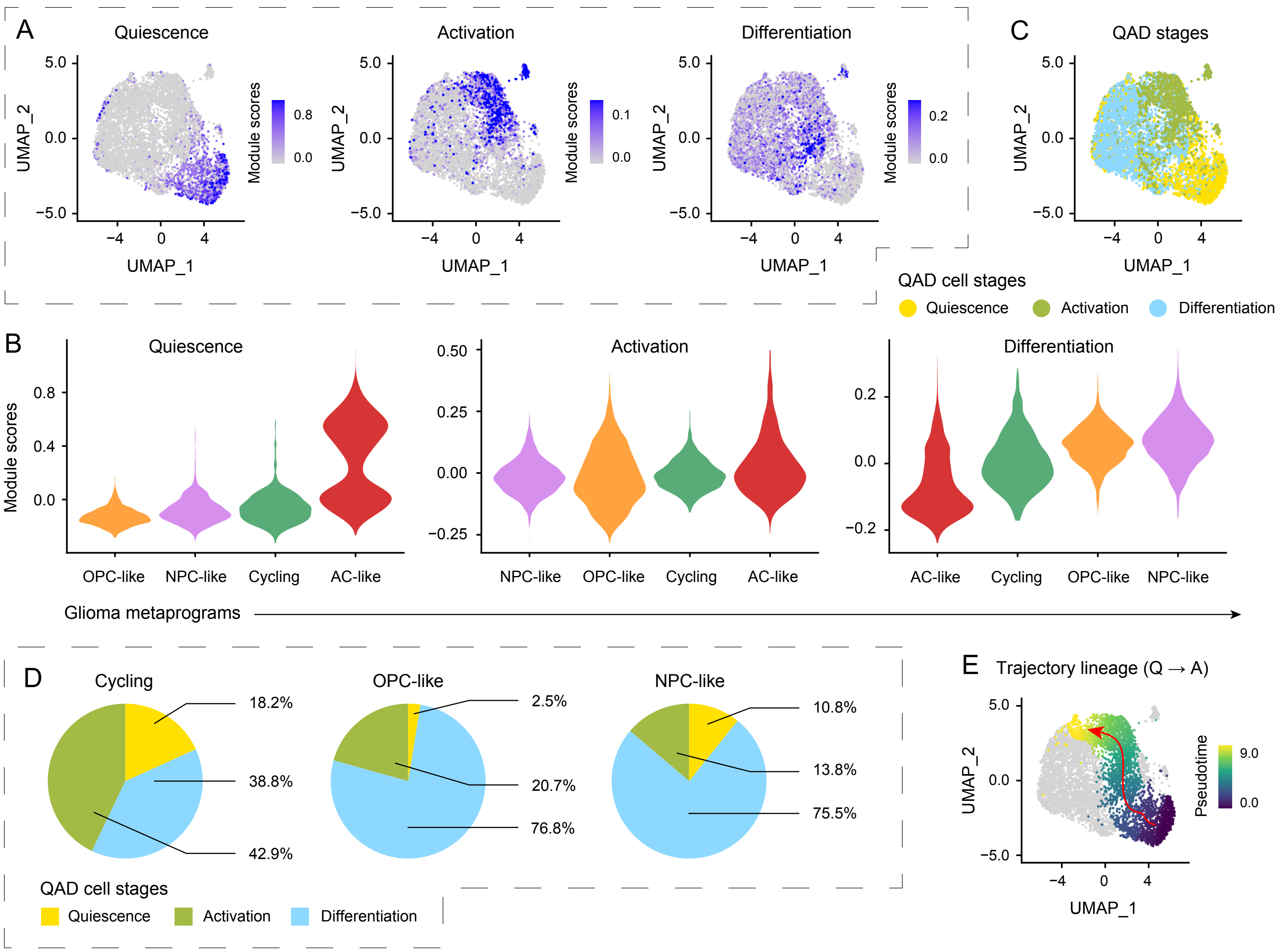

### Supplementary Figure 3.pdf

#### Supplementary Figure 3

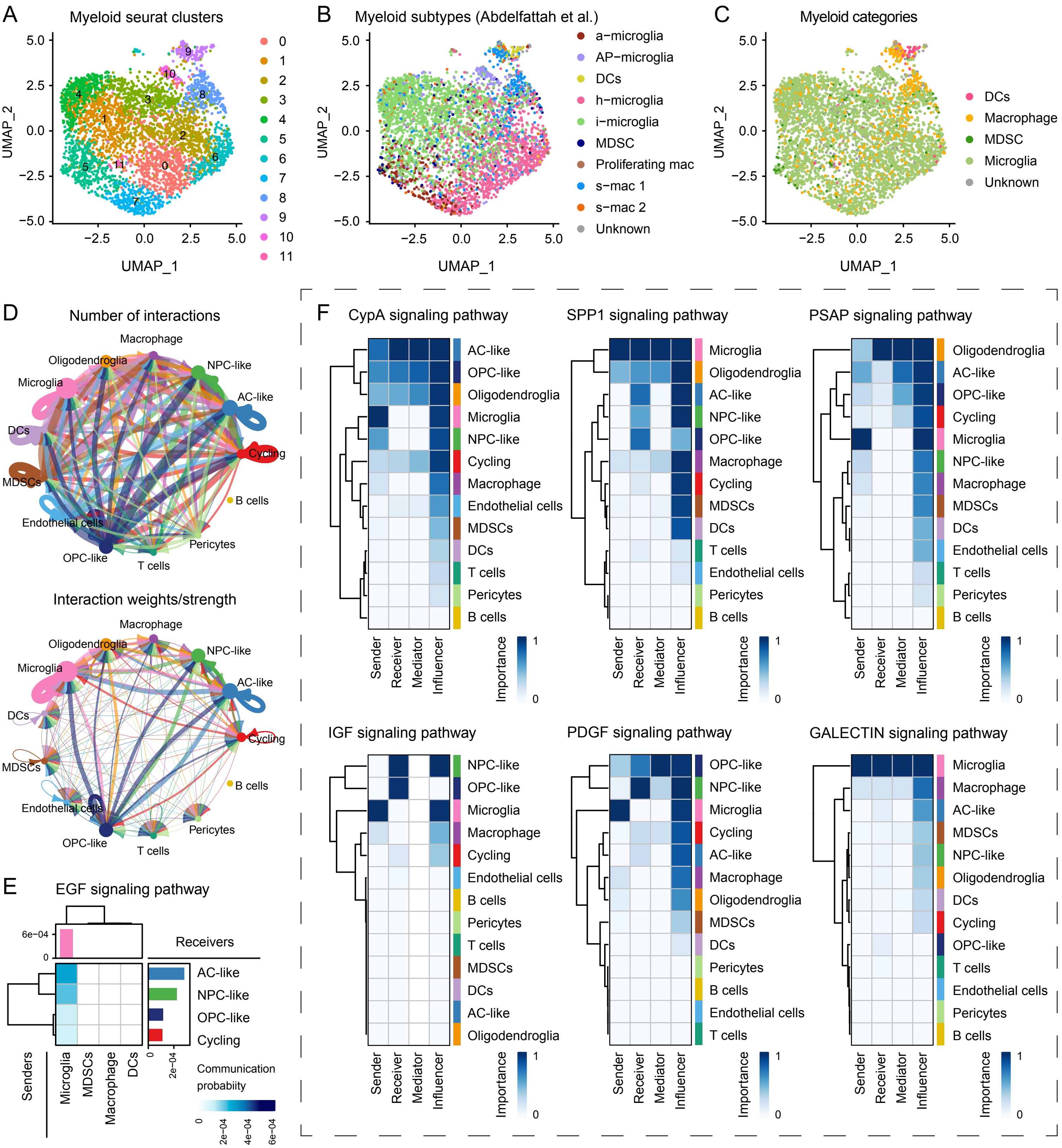

### Supplementary Figure 4.pdf

#### Supplementary Figure 4

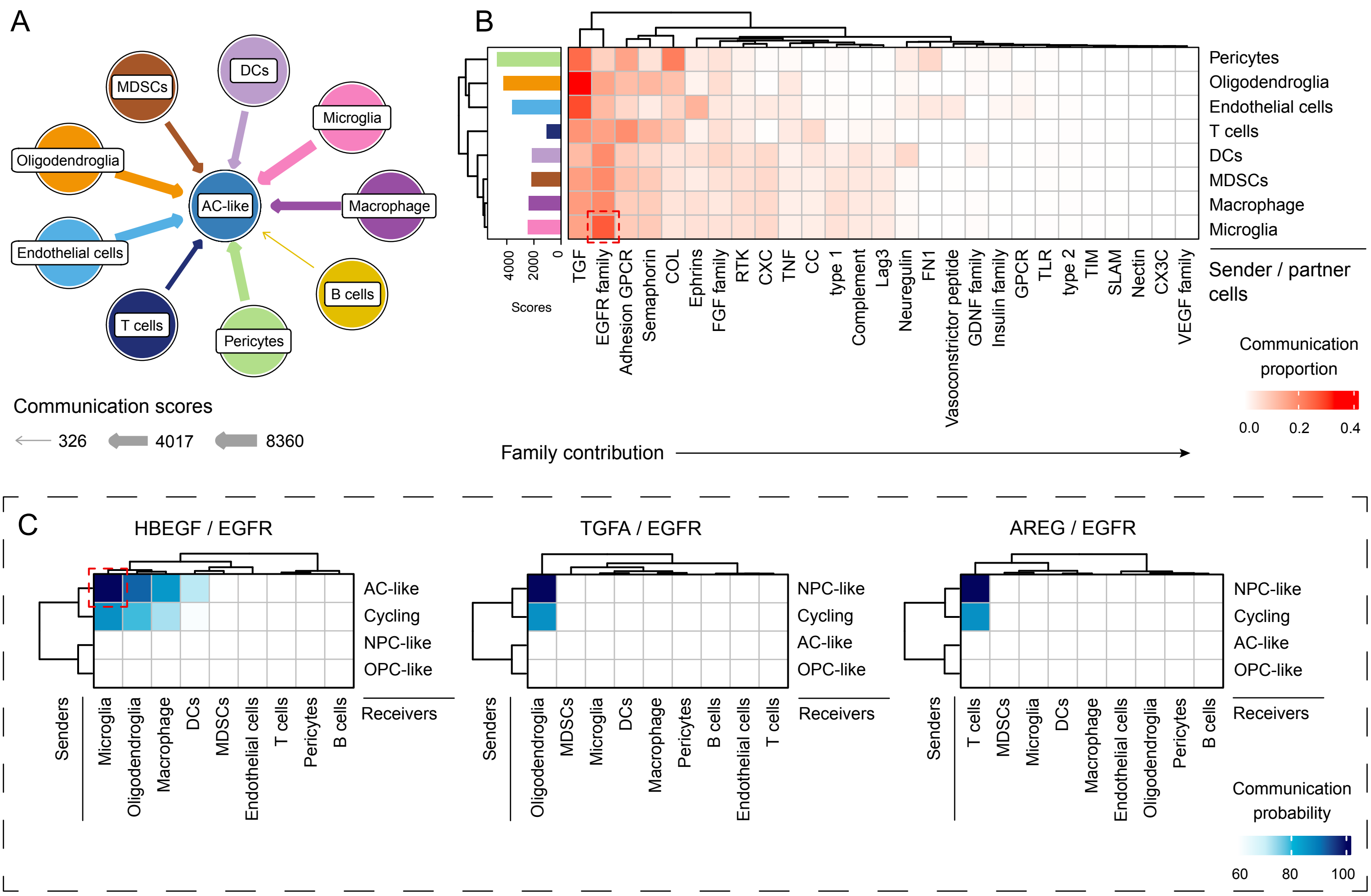

### Supplementary Figure 5.pdf

#### Supplementary Figure 5

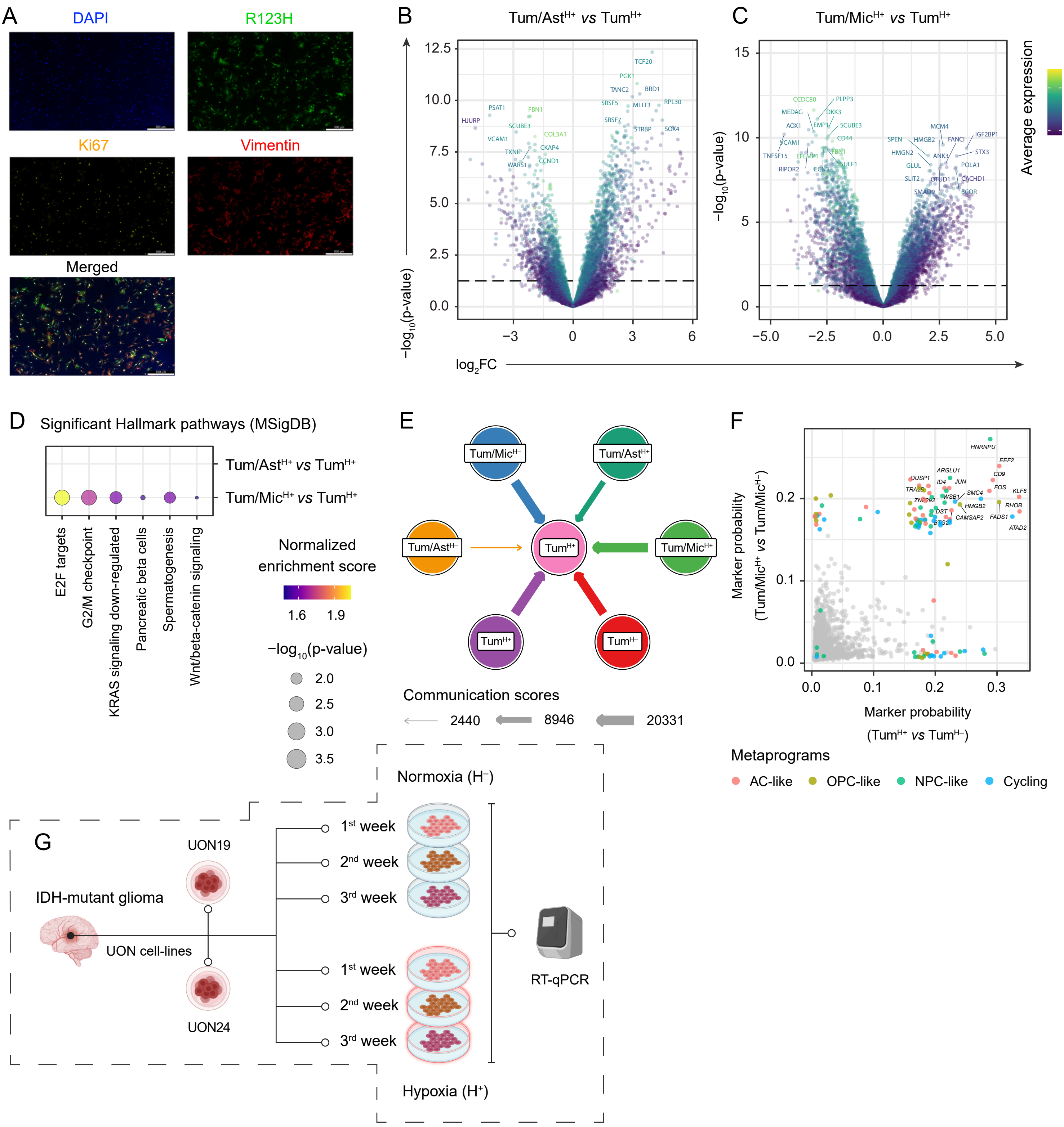

### Supplementary Figure 6.pdf

#### Supplementary Figure 6

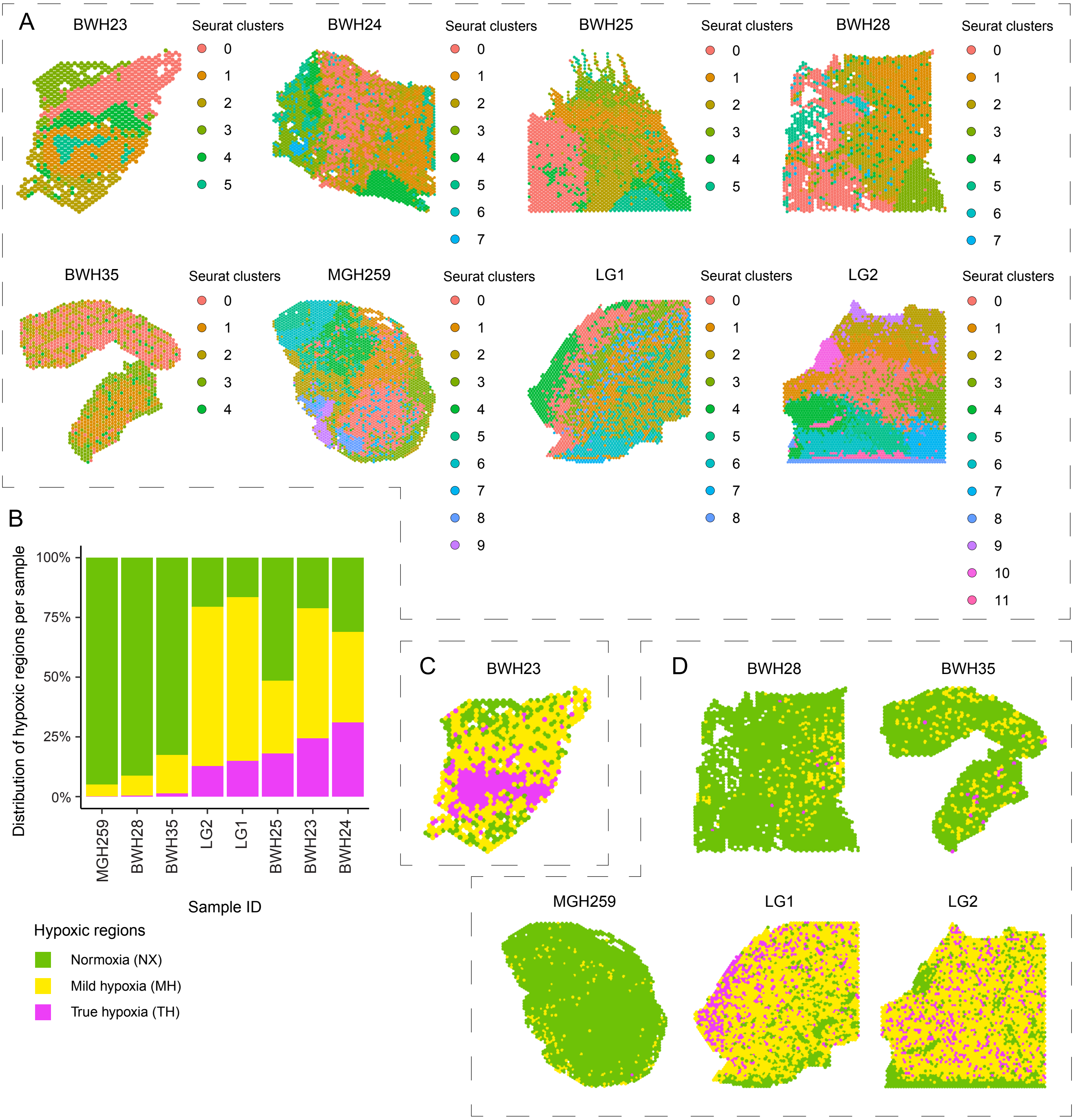

### Supplementary Figure 7.pdf

#### Supplementary Figure 7

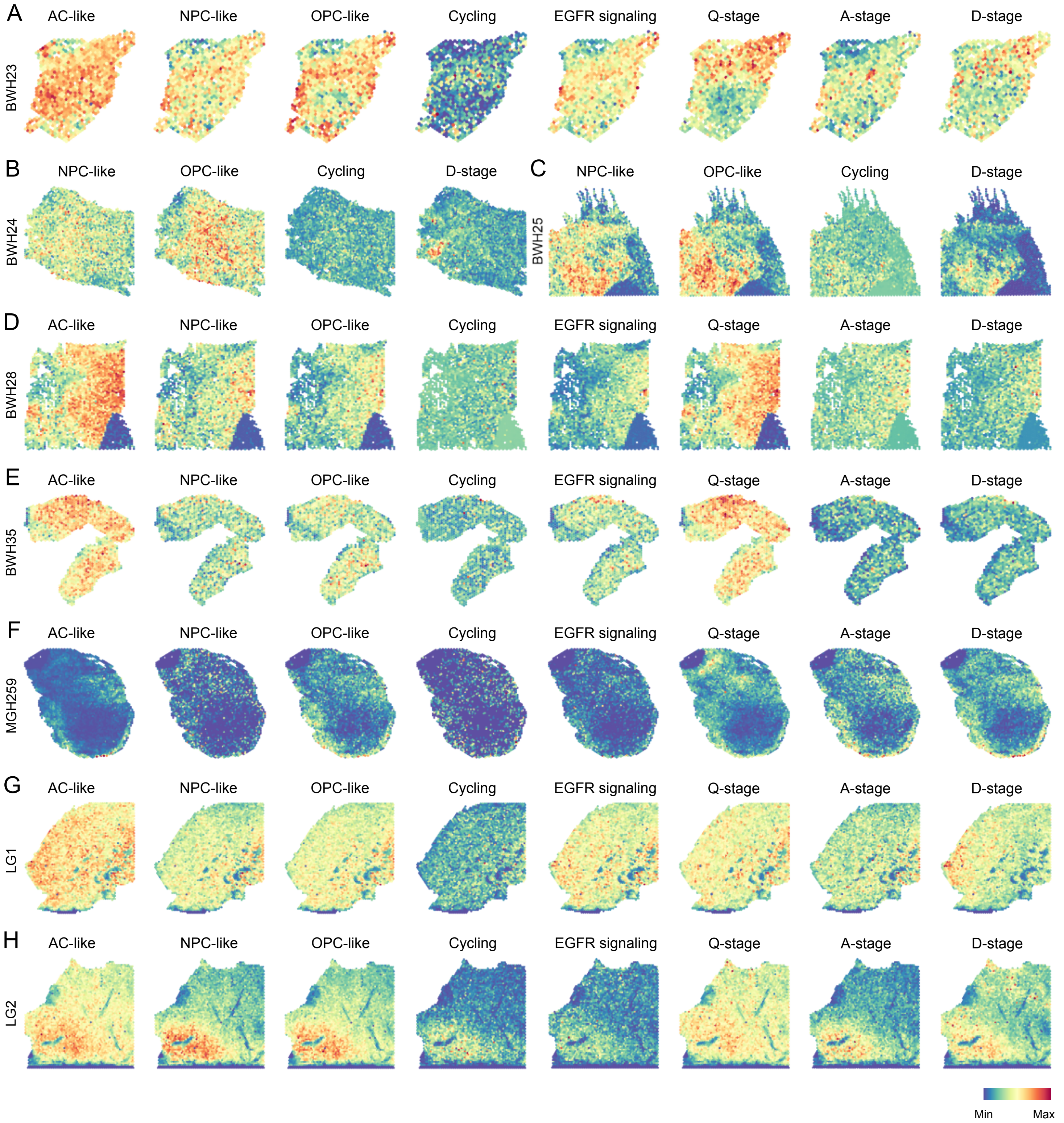

### Supplementary Figure 8.pdf

#### Supplementary Figure 8

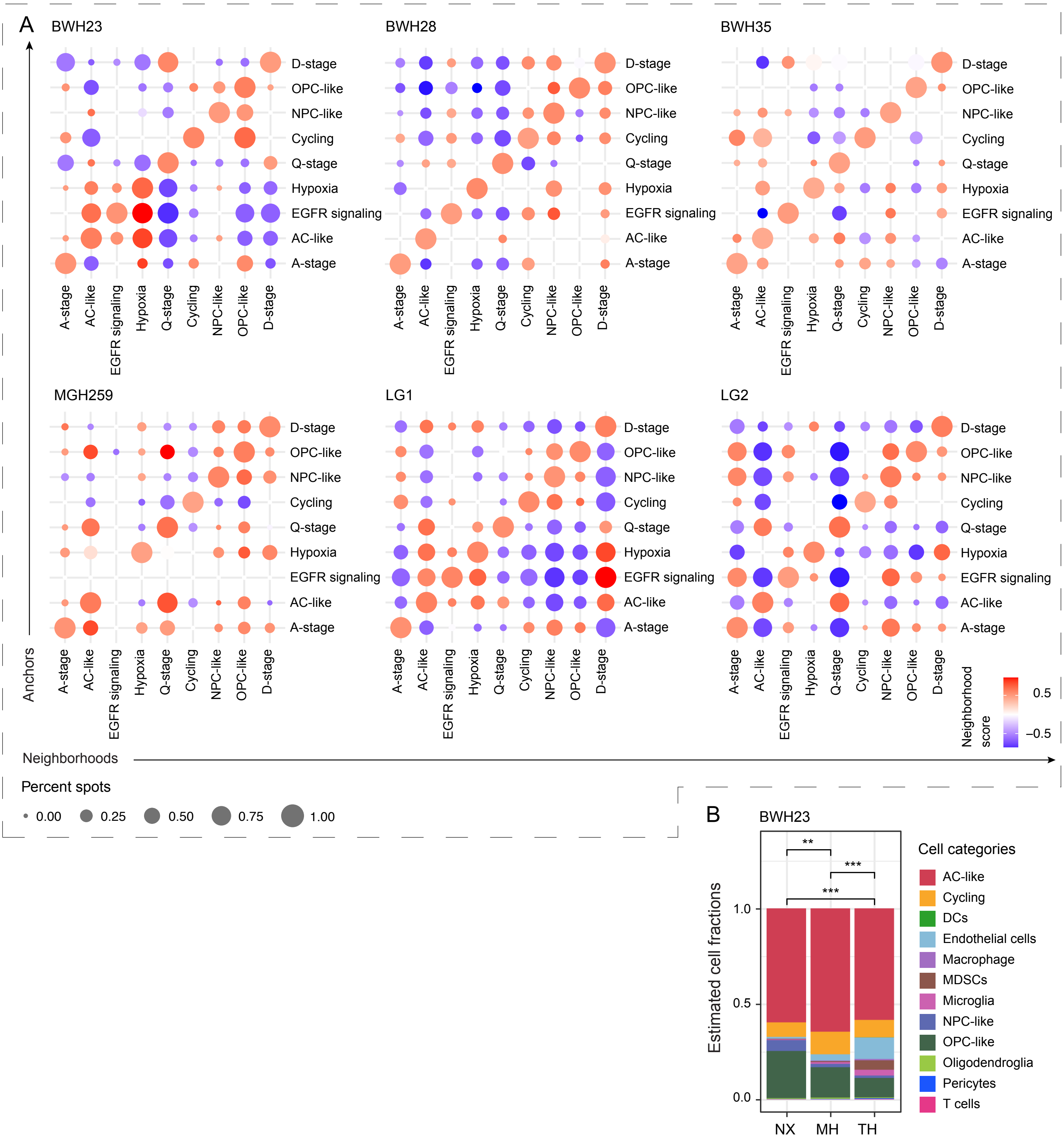

### Supplementary Figure 9.pdf

#### Supplementary Figure 9

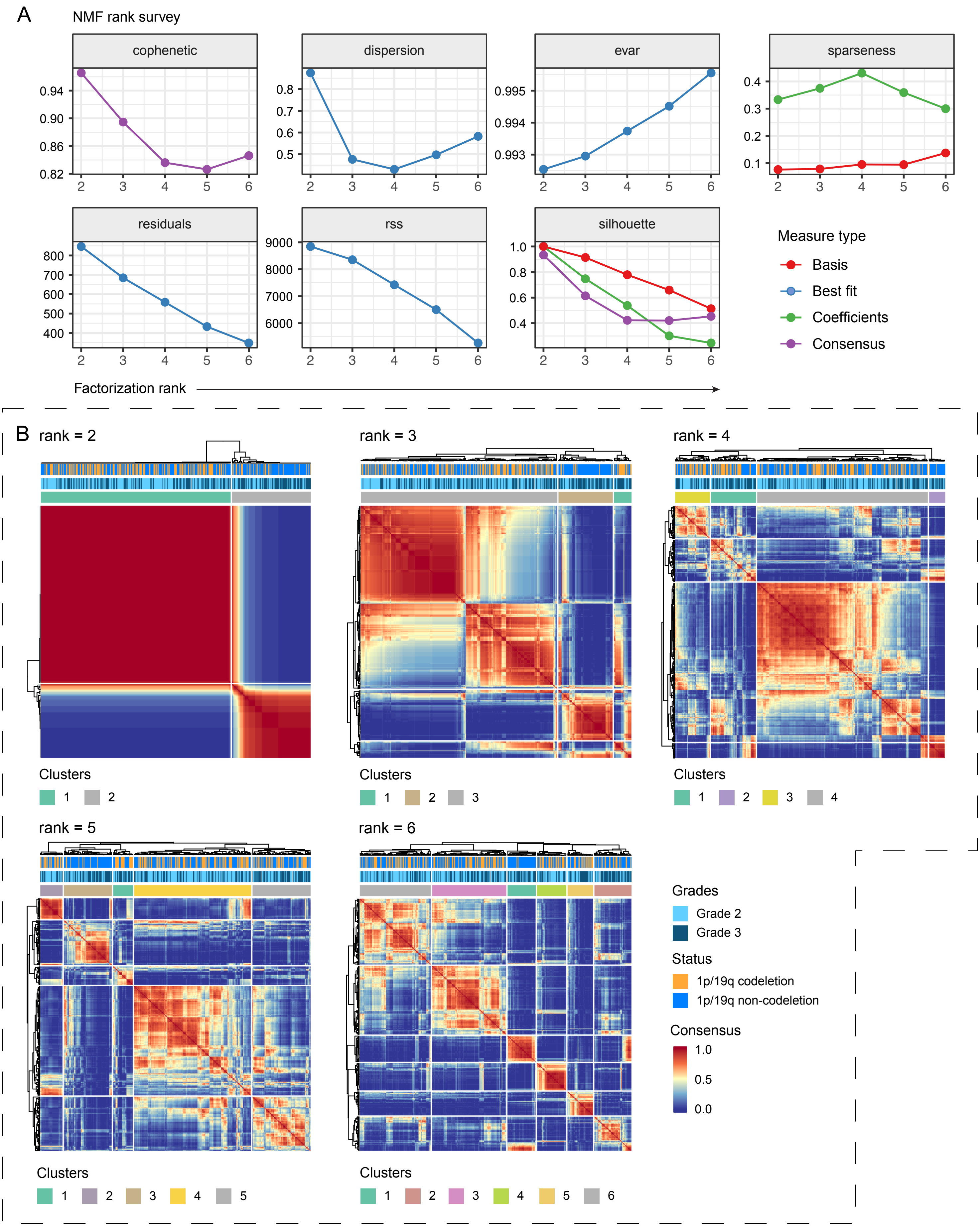

### Supplementary Figure 10.pdf

#### Supplementary Figure 10

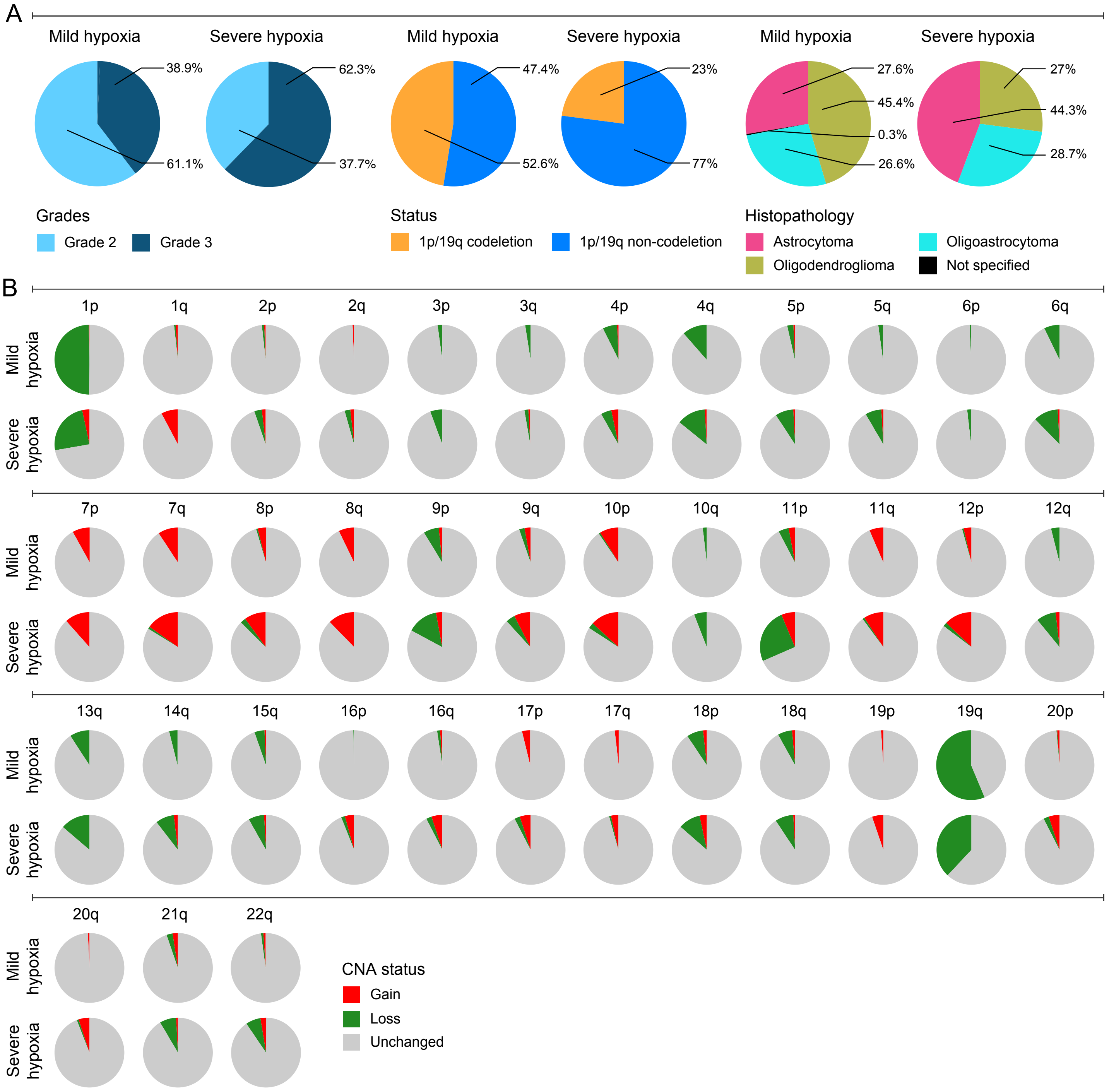

### Supplementary Figure 11.pdf

#### Supplementary Figure 11

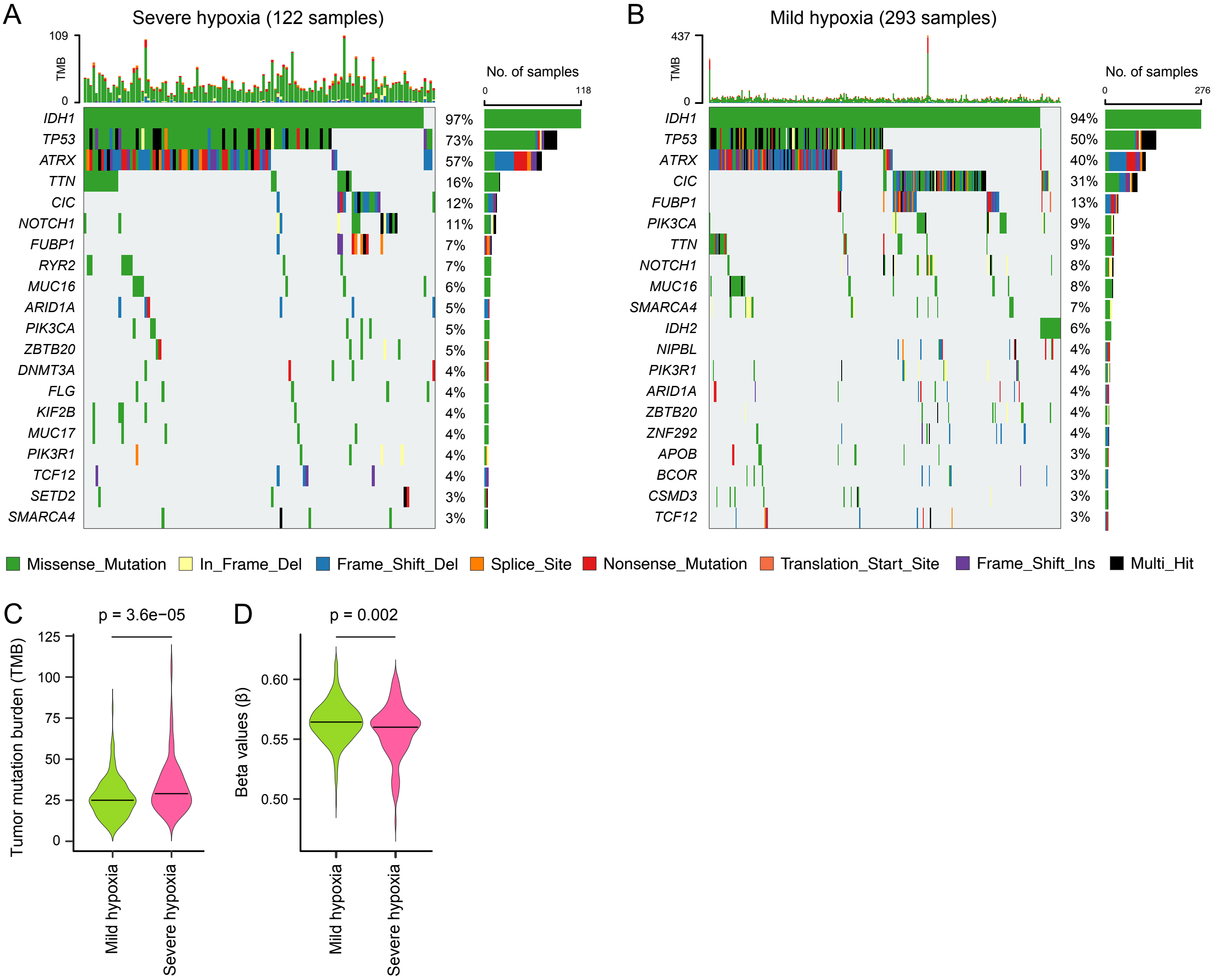

### Supplementary Figure 12.pdf

#### Supplementary Figure 12

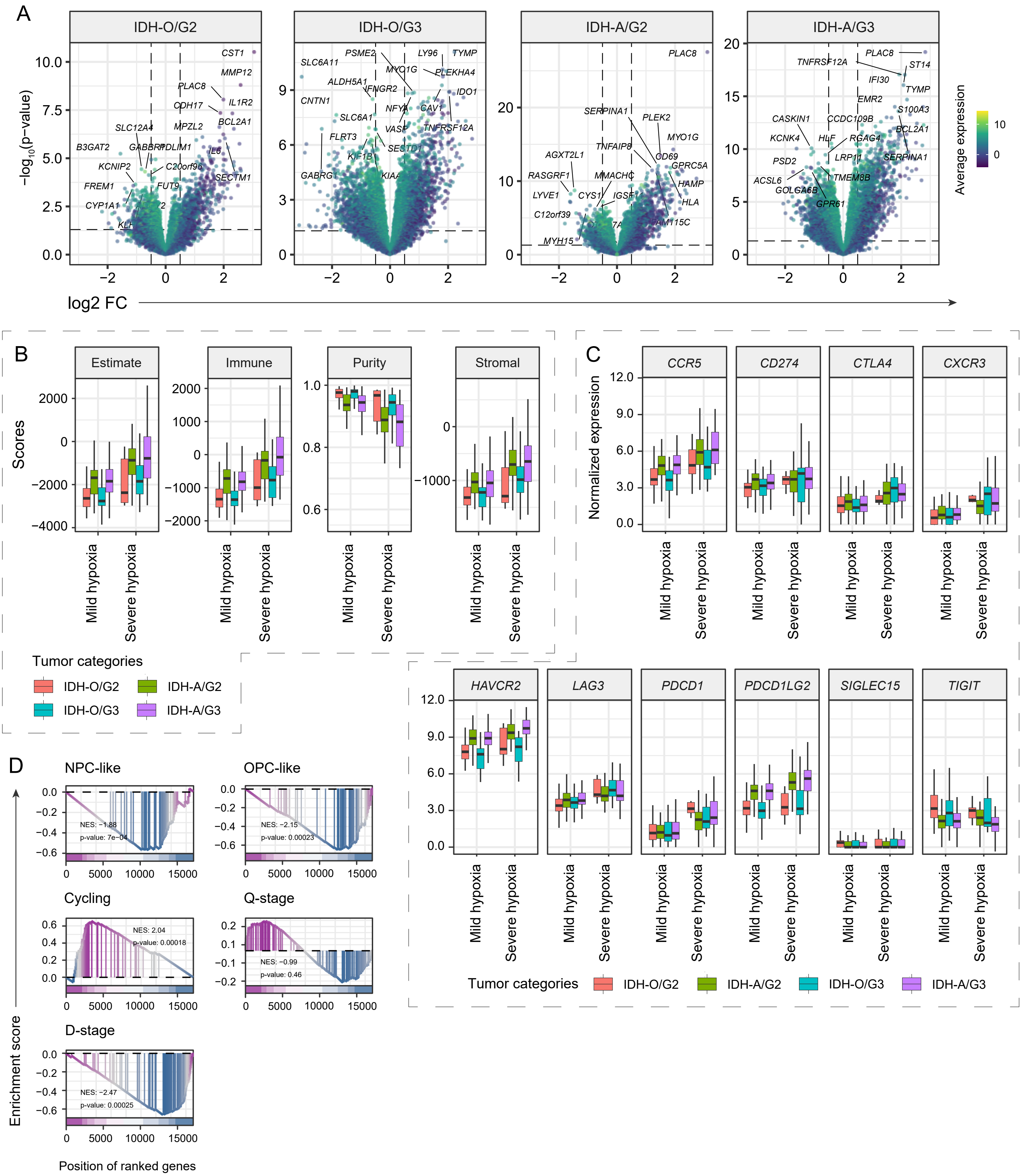

### Supplementary Figure 13.pdf

#### Supplementary Figure 13

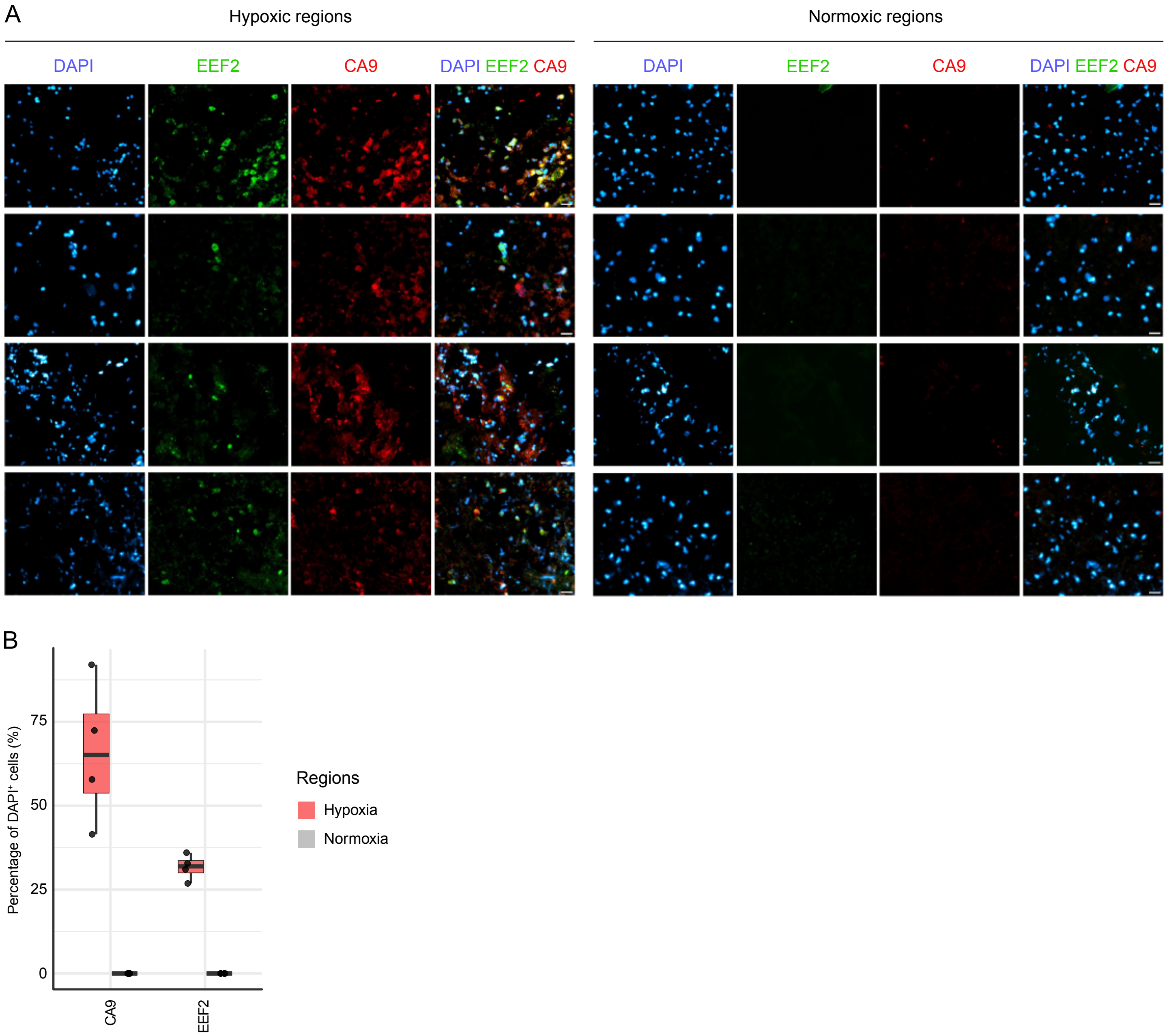
